# Copy number variants outperform SNPs to reveal genotype-temperature association in a marine species

**DOI:** 10.1101/2020.01.28.923490

**Authors:** Yann Dorant, Hugo Cayuela, Kyle Wellband, Martin Laporte, Quentin Rougemont, Claire Mérot, Eric Normandeau, Rémy Rochette, Louis Bernatchez

## Abstract

Copy number variants (CNVs) are a major component of genotypic and phenotypic variation in genomes. To date, our knowledge of genotypic variation and evolution has largely been acquired by means of single nucleotide polymorphism (SNPs) analyses. Until recently, the adaptive role of structural variants (SVs) and particularly that of CNVs has been overlooked in wild populations, partly due to their challenging identification. Here, we document the usefulness of Rapture, a derived reduced-representation shotgun sequencing approach, to detect and investigate copy number variants (CNVs) alongside SNPs in American lobster (*Homarus americanus*) populations. We conducted a comparative study to examine the potential role of SNPs and CNVs in local adaptation by sequencing 1,141 lobsters from 21 sampling sites within the southern Gulf of St. Lawrence, which experiences the highest yearly thermal variance of the Canadian marine coastal waters. Our results demonstrated that CNVs accounts for higher genetic differentiation than SNP markers. Contrary to SNPs, for which no significant genetic-environment association was found, 48 CNV candidates were significantly associated with the annual variance of sea surface temperature, leading to the genetic clustering of sampling locations despite their geographic separation. Altogether, we provide a strong empirical case that CNVs putatively contribute to local adaptation in marine species and unveil stronger spatial signal of population structure than SNPs. Our study provides the means to study CNVs in non-model species and highlights the importance of considering structural variants alongside SNPs to enhance our understanding of ecological and evolutionary processes shaping adaptive population structure.

## Introduction

The paradigm of local adaptation states that heterogeneous environmental conditions across the landscape can generate and/or maintain variation in morphology, physiology, behaviour and life history traits to maximize fitness under at least one set of habitat variables (Williams 1966; Savolainen et al. 2013; Hoban et al 2016). Therefore, investigating how genotypic variation is influenced by environmental heterogeneity is critical for understanding the evolution of adaptive traits. Local adaptation has been extensively studied in terrestrial landscapes and freshwater ecosystems (Salvolainen et al. 2013; Manel & Holderegger, 2013). While studies the field of marine ecological population genetics widely reported that local adaptation plays an important role in shaping the genetic structure of populations in marine habitats (e.g. Billerbeck et al., 2000; Pespeni & Palumbi 2013), much remains to be done toward deciphering the genomic basis underlying local adaptation in marine ecosystems (see reviews by Bernatchez 2016; Grummer et al. 2019; Palumbi et al., 2019). Marine species can represent an excellent model for studying adaptive genetic variation due to their generally large effective population size combined with high dispersal abilities resulting in high gene flow and minimal imprint of genetic drift (Palumbi 1992, Gagnaire et al., 2015, Tigano and Friesen 2016). Recent works pointed out that signal of local adaptation in marine species are often limited to small genomic regions, which identification specifically requires large genomic data set (Gagnaire et al. 2015).

Over the last decade, high throughput single nucleotide polymorphisms (i.e. SNPs) data have facilitated population genomic studies and substantially improved our understanding of the mechanisms underlying local adaptation (Savolainen et al. 2013). In particular, the development of reduced-representation shotgun sequencing approaches (i.e. RRS; including RADseq or GBS) has allowed genotyping of SNPs across large sample sizes to address the genetic basis of adaptation in non-model species, including marine organisms (Hess et al., 2013; Hemmer-Hansen et al., 2014; Guo et al., 2015, Bernatchez et al., 2019; Sandoval-Castillo 2018; Xuereb et al. 2018). Yet, to date, studies aiming to link genotypic variation with adaptation in wild populations have largely focussed on SNPs, which represent point mutation events,. In comparison, the role of structural variants has been overlooked due to technical and financial constraints (Wellenreuther et al., 2019, Mérot et al 2020). Structural variants (SVs) are genomic rearrangements affecting the presence, position or direction of a DNA sequence, including deletions, duplications, insertions, inversion or translocations (Mérot et al., 2020). SVs are widespread variants that shape genome architecture, and can cover a larger proportion of genetic variation than SNPs (Redon et al., 2006; Wellenreuther et al., 2019; Catanach et al., 2019). They can have functional consequences, notably impacting the regulation of gene expression (Gamazon & Stranger 2015) and recombination rate by diminishing the frequency of meiotic crossing-over, which can preserve the integrity of an adaptive haplotype (Rowan et al., 2019). SVs are increasingly recognized for their important role in a wide spectrum of evolutionary processes such as in environmental adaptation (Van’t Hof et al., 2016; Wellenreuther et al., 2017), reproductive isolation (Berdan et al. 2019; Laporte et al. 2019), life cycles and development (Mérot et al 2019; Cayuela et al., 2020), and speciation (Rieseberg 2001; Serrato-Capuchina & Matute 2018).

Copy number variants (CNVs) are a particular type of SV, which involves insertions, deletions and duplications of DNA sequences ranging in size from di-nucleotides repeats to millions of bases (Clop, Vidal & Amills, 2012). CNVs brings additional genomic diversity, that may complement information provided by SNPs concerning the extent of adaptive genomic divergence underlying inter-individual phenotypic variation (Nguyen et al., 2006, Levy et al., 2007). For example, while studies on human initially reported that the genomes of two unrelated individuals differ by ∼0.1% when considering only SNPs, the introduction of SVs as an additional source of variation revised this estimate up to ∼0.5% (Levy et al., 2007), with most of this difference due to CNVs (Redon et al. 2006; Levy et al. 2007). Although the majority of CNVs occurs within intergenic regions of eukaryote genomes (Emerson et al., 2008; Conrad et al., 2010), large CNVs may encompass entire genes (Collins et al., 2020), and modify downstream expression of regulatory regions as well as multiple protein-coding genes (i.e., pleiotropic effects; Gamazon and Stranger 2015). Originally associated with human genome studies (i.e. diseases, pathogen susceptibility, drug response, ancestry; Feuk, Carson & Scherer 2006; Wong et al., 2007; Ionita-Laza et al., 2009), CNVs have since been studied in a few other species, notably to understand domestication or speciation processes (Clop et al., 2012). So far, the role of CNVs in local adaptation remains poorly understood because studies in non-model species are just emerging (Wellenreuther et al., 2019). Moreover, they are often restricted to one or a few large-effect CNVs or duplicated genes (Tigano et al., 2018, Smith et al., 2017; Rinker et al., 2019; Nelson et al., 2018, Prunier et al., 2019), without testing more systematically the contribution of all detected CNVs relatively to SNPs (Mérot et al 2020). Also, CNVs have rarely been studied on a large subset of individuals due to financial and technical constraints because classical approaches to detect CNVs, such as paired-end mapping, split read, *de novo* assembly or depth of coverage analysis, require a reference genome and whole-genome resequencing at deep coverage (Roca et al., 2019). However, recent technological and methodological advances in sequence analyses hold promise to address such issues. In particular, an approach has recently been proposed to use cost-efficient reduced-representation sequencing data to distinguish multiple-copy (e.g. duplicated, CNV) from non-duplicated single-copy loci (e.g. McKinney et al., 2017a). Originally developed to deal with specific issues on whole genome duplications and mixed ploidy problems in salmonids, McKinney’s et al. (2017a) method has been used to filter SNPs and to avoid miscalled genotypes, but it can also inform on CNVs.

Here, we conducted a study in the American lobster (*Homarus americanus*) to compare the association of SNPs and CNVs to thermal conditions in the southern Gulf of St. Lawrence, Canada, with the broader goal of increasing our understanding of the potential role of CNVs in local adaptation in non-model marine species. This species appears to be a relevant model for investigating this issue as recent insights from crustacean genome assemblies indicate that repeat regions represent a prominent proportion of the genome (Song et al., 2016; Zhang et al 2019). While past studies on the American lobster genome showed evidence of diploidy and estimated a genome size roughly 4.5 Gb (Jimenez et al., 2010), recent studies in other crustacean genomes suggested that life cycles strategies and environment could be major determinants in genome evolution, notably by promoting the expansion of its size (Alfsnes, Leinaas and Hessen, 2017). We thus hypothesized that CNVs could play an adaptive role in *H. americanus*. Based on previous genome-wide SNP studies in the same system (Benestan et al., 2015, Dorant et al., 2019), we anticipated a weak level of genetic differentiation among populations. Conversely, whereas the average mutation rate at CNV loci is generally reported to be higher than SNPs, various studies also reported that CNV evolution can be induced by the environment and then contribute to inter-population differences (Kofler, Nolte and Schlötterer, 2015; Sudmant et al., 2015). Hence, given the different properties of CNV and SNP markers, we predicted distinct patterns of population genetic divergence between these two types of genetic variants. Furthermore, as high gene flow may swamp beneficial alleles and potentially hamper local adaptation in marine species (Tigano & Friesen 2016), we predicted that CNVs could provide an alternative molecular basis for adaptation to temperature in the American lobster.

First, we combined reduce-representation sequencing and sequence capture enrichment (so-called Rapture; Ali et al., 2016) to sequence 1,141 lobsters and genotype CNVs and SNPs following the approach proposed by McKinney et al. (2017a). Second, we compared the extent of genetic differentiation among sampling sites using the two types of markers. Third, we examined the potential role of CNVs and SNPs in local adaptation by investigating their respective associations with temperature. Overall, our study highlights the importance of CNVs as a major source of genetic polymorphism in the genome of the American lobster which may contribute to its adaptation to local thermal conditions, and as such, our understanding of the species’ population biology.

## 2. Material and Methods

### 2.1 Sample collection and DNA extraction

A total of 1,141 lobster samples were collected from 21 sites between May and July in 2016 in the southern Gulf of St.Lawrence, Canada (figure 1a and Table S1). DNA was extracted from the distal half of a walking leg of each lobster using salt extraction (Aljanabi & Martinez, 1997) with an additional RNase treatment following the manufacturer’s instructions (Qiagen Inc., Toronto, Ontario, Canada). DNA quality was assessed using 1% agarose gel electrophoresis. Genomic DNA concentrations were normalized to 20 ng/µl based on a fluorescence quantification method (AccuClear™ Ultra High Sensitivity dsDNA Quantitation Solution). Individual reduced-representation shotgun sequencing (i.e. RRS) libraries were prepared following the Rapture approach (Ali et al., 2016). This approach is a form of RRS sequencing, which combines double-digested libraries (i.e. GBS or RADseq) with a sequence capture step. Obtaining appropriate depth of coverage through traditional RADseq approaches is often very costly, particularly so for large sampling design in terms of number of locations and individuals, and/or for species with a large genome size. The sequence capture step allows targetting a panel of informative and high quality RAD loci, which have been discovered and selected from an initial RADseq experiment. As such, these captured loci still represent a sampling of the genome-wide variation. Rapture thus represents a cost-effective and flexible approach which allowed sequencing a large number of samples with a high sequencing depth, thereby enabling efficient generation of a large population genomic dataset. More specifically, we used the same 9,818 targeted loci previously used for the American lobster and all the details about the wet protocol are described in Dorant et al. (2019). All Rapture libraries were sequenced on the Ion Torrent p1v3 chip at the Plateforme d’analyses génomiques of the Institute of Integrative and Systems Biology (IBIS, Université Laval, Québec, Canada http://www.ibis.ulaval.ca/en/home/). Two rounds of sequencing (i.e. two separated chips) were conducted for all Rapture libraries.

**Figure 1:**
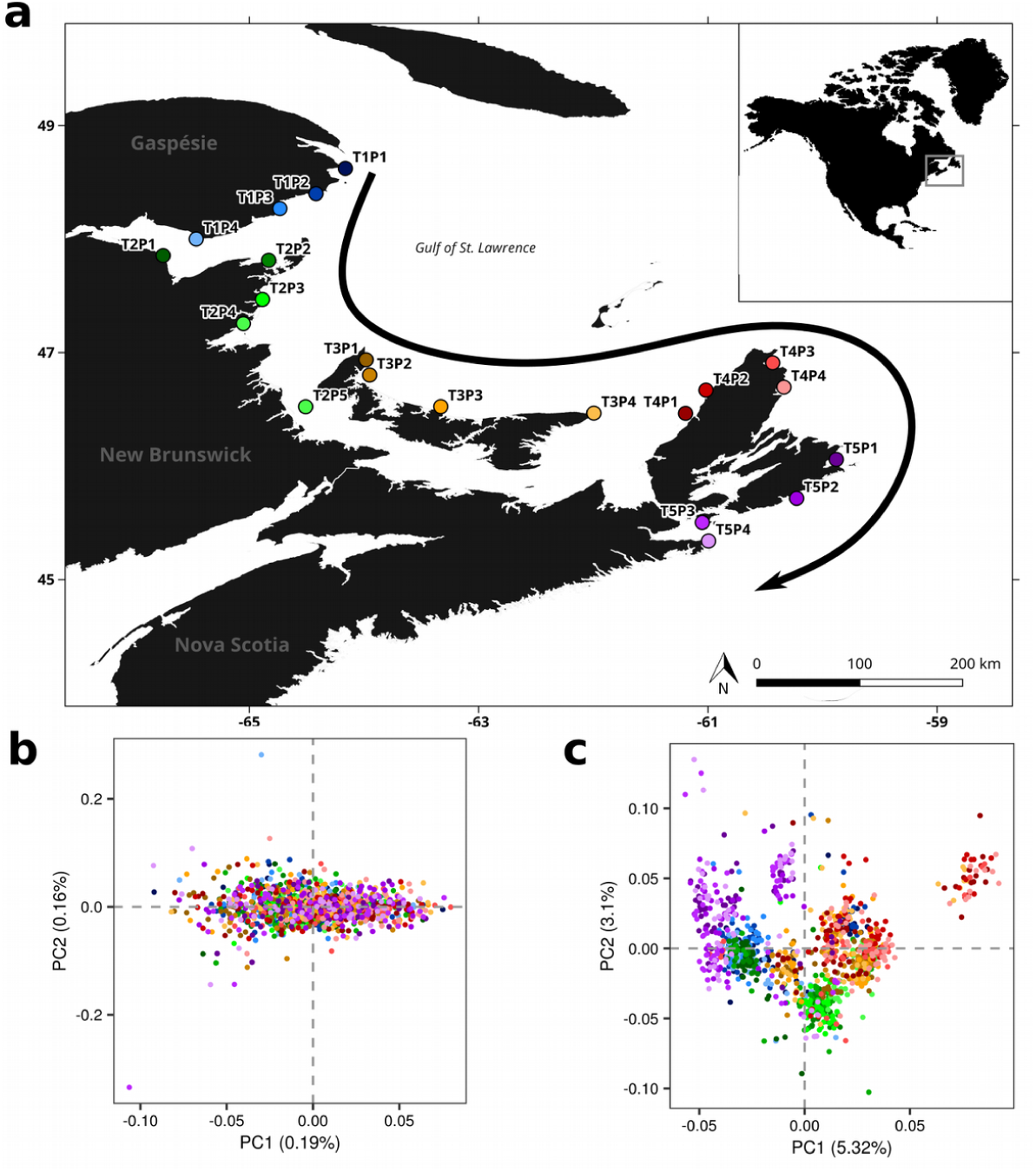
Geographic distribution and genetic structure of American lobster in southern Gulf of St. Lawrence. **a**. Map of American lobster sample collection. Sampling sites are represented by colored circles. The black arrow represents the direction of the major current pattern within the region during the larval drift period between July and September (Galbraith et al., 2015). **b**. PCA plot inferred from the 13,854 SNPs genotyped. **c**. PCA plot inferred from normalized read-depth of 1,521 CNV loci identified. PCAs display the individuals colored according to the same color code depicted in the sampling map (1a).

### 2.2 Genotyping and data filtering

Single-end raw reads were trimmed to 80 bp and shorter reads were removed using *cutadapt* (Martin, 2011). We only retained samples with at least 200,000 sequencing reads for downstream pipeline. This threshold was fixed to remove poorly sequenced samples and maximize the expected locus read depth across samples. *BWA mem* (Li 2013) was used to map the resulting individual-based sequences to the Rapture reference catalog previously developed by Dorant et al. (2019) that consisted of 9,818 targeted sequences in total. SNP discovery was done with *Stacks* v.1.48 (Catchen, Hohenlohe and Bassham, 2013). A minimum stack depth of four (*pstacks* module : m=4) and a maximum of three nucleotide mismatches (*cstacks* module : n=3) were allowed. We ran the *population* module requiring a locus to be genotyped at a frequency greater than 60% in at least four out of 21 sampling sites. We then filtered genotype data and characterized CNV loci using filtering procedures and custom scripts available in *stacks_workflow* (https://github.com/enormandeau/stacks_workflow). In an initial step, we filtered the raw VCF file keeping only genotypes that i) showed a minimum depth of four (parameter “m” hereafter), ii) were called in at least 70% of the samples in each site (parameter “p” hereafter), and iii) that had an MAS value of two (parameter “S” hereafter), the *05_filter_vcf_fast*.*py* available in *stacks_workflow*. The MAS parameter, which is akin to the minor allele frequency (MAF) or minor allele count (MAC) filter, refers to the number of different samples possessing the minor allele. Here, we implemented the MAS filter for short-read data as it does not suffer from the same biases inherent to MAF and MAC, which are boosted by genotyping errors where one heterozygous sample is erroneously genotyped as a rare-allele homozygote. Then, we removed samples showing more than 15% of missing data. To control for putative sample DNA contamination (e.g. which can occur during DNA normalization, library preparation), relatedness between samples and the inbreeding coefficient were estimated. Relatedness was estimated following the equation proposed by Yang et al. (2010) and implemented in *vcftools*. Constitutively, the relatedness parameter expects that unrelated individuals tend to zero, while a value of one is expected for an individual with itself (Yang et al., 2010). A relatedness coefficient around 0.5 should represent siblings and high value of relatedness between two different individuals may represent identical twins or clones, which is not expected in this species. Hence, in cases where pairs of samples showed a relatedness coefficient > 0.90, we excluded the one sample (out of the two being compared) exhibiting the higher level of missing data. The inbreeding coefficient (*F*_*IS*_) was estimated for each sample using a method of moments implemented in *vcftools*. From graphical observation of sample inbreeding, we defined a cutoff value (i.e. -0.25) to exclude outlier samples. We then removed all samples exhibiting putative DNA contamination from the raw VCF file and then re-ran the *05_filter_vcf_fast*.*py* from *stack_workflow*, keeping the same parameters previously used (i.e. m=4; p=70; S=2).

### 2.3 Identification of copy number variants

To explore locus duplication and then identify putative CNVs, we based our analyses on the previous work by McKinney et al (2017a). Using population genetic simulations and empirical data, McKinney et al. (2017a) demonstrated that a set of simple summary statistics (e.g. the proportion of heterozygous and read ratio deviation of alleles) could be used to confidently discriminate SNPs exhibiting a duplication pattern without the need for a reference genome. Thus, we investigated SNP “anomalies” based on a suit of four parameters to discriminate high confidence SNPs (hereafter singleton SNPs) from duplicated SNPs: (1) median of allele ratio in heterozygotes (MedRatio), (2) proportion of heterozygotes (PropHet), (3) proportion of rare homozygotes (PropHomRare), and (4) inbreeding coefficient (*F*_*IS*_). Each parameter was calculated from the filtered VCF file using the *08_extract_snp_duplication_info*.*py* available in *stacks_workflow*. The four parameters calculated for each locus were plotted against each other to visualize their distribution across all loci. Based on the methodology of McKinney et al (2017a), and by plotting different combinations of each parameter, we graphically fixed cut-offs for each parameter. Full details of this step are available in table S2 and figure S1. Finally, two separate data sets were generated: the “SNP dataset”, based on SNP singletons only, and the “CNV dataset”, based on duplicated SNPs only. To construct the SNP dataset, we kept the genotype calls from the VCF file containing singleton SNPs, and post-filtered these by keeping all unlinked SNPs within each locus using the *11_extract_unlinked_snps*.*py* available in *stacks_workflow*. Briefly, the first SNP is kept and all remaining SNPs showing strong genotype correlation are pruned (i.e. two SNPs show strong genotype correlation if samples with the minor allele in one of the SNPs have the same genotypes as samples with the minor allele in the other SNP more than 50% of the time). The procedure was repeated until all SNPs were either kept or pruned. To construct the CNV dataset, we extracted the locus read depth of SNPs identified as duplicated using *vcftools*. Note that the CNV dataset contains duplicated loci that could be invariant in copy number among the 21 sampling sites. Hence, for simplicity, we use the term “CNV” to represent all loci classified as duplicated by our approach. As libraries sequenced at a greater depth will result in higher overall read counts, CNV locus read counts were normalized to account for differences in sequencing coverage across all samples. Normalization was performed using the trimmed mean of M-values method originally described for RNA-seq count normalization and implemented in the R package *edgeR* (Robinson & Oshlack, 2010). The correction accounts for the fact that for an individual with a higher copy number at a given locus, that locus will contribute proportionally more to the sequencing library than it will for an individual with lower copy number at that locus. Finally, the resulting CNV dataset was a matrix of normalized read count for each individual at each CNV locus.

### 2.4 Genetic differentiation analyses

We estimated pairwise *F*_*ST*_ (Weir & Cockerham, 1984) values for the SNP dataset using *stamppFst* function from the R package *StAMPP* v.1.5.1 (Pembleton et al., 2013). To estimate population genetic differentiation of loci identified as CNVs, we calculated the variant fixation index *V*_*ST*_ (Redon et al. 2006). *V*_*ST*_ is an analog of *F*_*ST*_ estimator of population differentiation (Weir & Cockerham, 1984) and is commonly used to identify differentiated CNV profiles between populations (Redon et al., 2006; Dennis et al., 2017; Rinker et al 2019). For each pairwise population comparison, *V*_*ST*_ was estimated considering (V_T_-V_S_)/V_T_, where V_T_ is the variance of normalized read depths among all individuals from the two populations and V_S_ is the average of the variance within each population, weighed for population size (Redon et al 2006). Then, we compared the magnitude of population differentiation estimated from SNPs and CNVs, respectively. Finally, we performed principal components analyses (PCAs) to visualize the pattern of genetic differentiation based on either all SNPs or all CNVs.

### 2.5 Genotype-Environment Association Analysis

Marine climatic data were extracted from geoTiff layers of the *MARSPEC* public database [http://marspec.weebly.com/modern-data.html] (Sbrocco & Barber, 2013). Five environmental layers (30 arcseconds resolution, i.e. ∼1km) related to Sea Surface Temperature (hereafter SST) were considered. These include the annual mean, annual range, annual variance, the minimum value observed for the coldest month and the maximum value observed for the warmest month. We focused on sea surface temperature, since this environmental parameter is one of the most critical for larval deposition and survival for the American lobster (Quinn et al., 2013, Quinn et al., 2015). Furthermore, lobsters used in this study were collected by Canadian fishers close to the shoreline, within < 20m depth, where sea surface temperature is a reasonable proxy of temperature in the entire water column. For all five temperature layers, we defined a circular buffer of 15km radius around each of the 21 sampling sites and values within this buffer were averaged to minimize pixel anomalies due to known biases of correction algorithms used for remote sensing, especially in coastal areas (Smit et al 2013). Collinearity between explanatory environmental variables was checked using the Pearson correlation index as well as a Draftsman’s plot. Among variables exhibiting high level of correlation (i.e. r > 0.6), we kept only one of them based on its biological relevance (see explanations in Results).

We performed a redundancy analysis (RDA) to investigate the association between environmental variables and genetic variation (either SNP or CNV data sets) using the *vegan* R library (Oksanen et al., 2018) and following Forester et al. (2018) (see https://popgen.nescent.org/2018-03-27_RDA_GEA.html for details). We applied a forward stepwise selection process to environmental predictors via the *OrdiR2step* function (1,000 permutations; limits of permutation *P* < 0.05) in order to identify environmental variables that significantly explained overall genetic variation (i.e. maximising adjusted R^2^ at every step). Ultimately, global and marginal analyses of variance (ANOVA) with 1,000 permutations were performed to assess the significance of the models and evaluate the contribution of each environmental variable. Once genetic markers were loaded against the RDA axes, candidates for positive response with environmental predictors were determined to be those that exhibit more than 2.5 standard deviations (SD) away from the mean (*P* < 0.012). More precisely, this cut-off represents ±2.5 SD from the mean loading of each axis, where outlier SNPs are identified from a two-tailed normal distribution and the mean is centered to 0 (see https://popgen.nescent.org/2018-03-27_RDA_GEA.html and Forester et al. 2018 for more details).

We also used the Latent Factor Mixed Models implemented in the R library *LFMM 2* (Caye et al,. 2019) for SNPs and linear mixed-effects models (LME) implemented in the *lme4* R library (Bates et al., 2015) for CNV data. This second analysis aimed to document the form and strength of the relationship between genetic markers and environmental predictors. In the LME model, sampling sites were selected as a random effect and environment parameters as the explanatory variables. Then, the genetic information (i.e SNP genotypes for singleton loci and normalized read depth for the putative CNV loci) were introduced in the model as the response variable. The effect of temperature predictors was assessed using ANOVA F-test applied on regression model output. To account for multiple testing issues, we used a false discovery rate (FDR) following the method of Benjamini & Hochberg (1995) for both LFMM and LME results. The p-value was adjusted to control the threshold for statistical significance in multiple comparisons at α = 0.01. To minimize potential false positives among candidate markers, we defined the set of best candidate loci for local (thermal) adaptation as those that were found to be associated with at least one environmental factor in both RDA and statistical regression models (i.e. LFMM and LME).

The relationship between CNV read depth (normalized) and the environmental predictors displayed a non-linear growth-like model for most candidates CNVs (see figure S4). Hence, we evaluated the fit of Gompertz (i.e. four parameters) and logistic functions (i.e. four to five parameters) for each locus. Model selection was done using the function *mselect* from the R library *drc* v3.0-1 (Ritz, 2015), and the model exhibiting the smallest log-likelihood coefficient was assumed to have the best fit to the data. Additionally, we assessed the clustering pattern among the 21 sampling sites using a Principal Component Analysis (PCA) on the candidate CNVs associated with the environmental predictors. The number of “meaningful” principal component (PCs) axes to retain for interpretation and downstream analyses was assessed based on the broken stick distribution (Legendre and Legendre, 2012).

### 2.6. Defining discrete copy number categories using clustering approach

For each CNV locus putatively associated with environmental variable, we examined whether discrete copy number categories could be drawn from the distributions of normalized read depth using a model-based unsupervised clustering approach implemented in the R library *Mclust*, V. 5.4.4 (Fraley et al., 2012). *Mclust* uses finite mixture estimation via iterative expectation maximization steps (EM) for a range of k components and the best model is selected using the Bayesian Information Criterion (BIC). For each locus, individuals were classified in discrete clusters read depth (that we coin copy number groups) that likely reflect variation in copy number of any given locus. Based on the copy number group characterizing each individual for a given locus, we were then able to assess the geographic distribution of copy number groups across all 21 sampling sites.

## 3. Results

### 3.1 Data processing and classifying singleton and CNVs loci

Rapture sequencing yielded an average of 431,350 (SD = 172,538) reads per sample before any quality filtering. From the 1,141 samples sequenced, 60 (5%) were removed based on individual’s quality filtering (i.e. 28, 25, 7 and 0 individuals for low sequencing depth threshold, up to 15% missing data, abnormal pattern of heterozygosity and abnormal relatedness, respectively). From the 1,081 remaining samples, the median number of reads per individual calculated for each sampling sites ranged from 364,437 ± 86,385 reads for T4P4 to 561,877 ± 219,624 reads for T3P1 (table S1). After the first SNP calling process, 26,005 SNPs spread over 6,946 loci of 80 bp were successfully genotyped in at least 70% of the individuals; these SNPs formed a pre-cleaned VCF file prior to the identification of singleton vs. duplicated SNPs. Overall, we observed that the proportion of missing data in this pre-filtered dataset was limited (i.e. 3.5% missing data overall). The characterization of SNPs returned 14,534 SNPs as singletons and 9,659 SNPs as duplicated (figure 2; see table S3 and figure S2 for details). The final SNPs singleton data set contained 13,854 SNPs, after keeping only unlinked SNPs within loci spread over 5,362 loci of 80 bp. The average sequencing depth among these 13,854 SNPs was 28X ranging from 6.2X to 126.7X. Finally, the CNV dataset consisted of read depth information extracted from the 1,521 loci, which comprises the 9,659 SNPs classified as duplicated. The average sequencing depth among these 1,521 CNV loci was 45.6X ranging from 7.4X to 2,560X.

**Figure 2:**
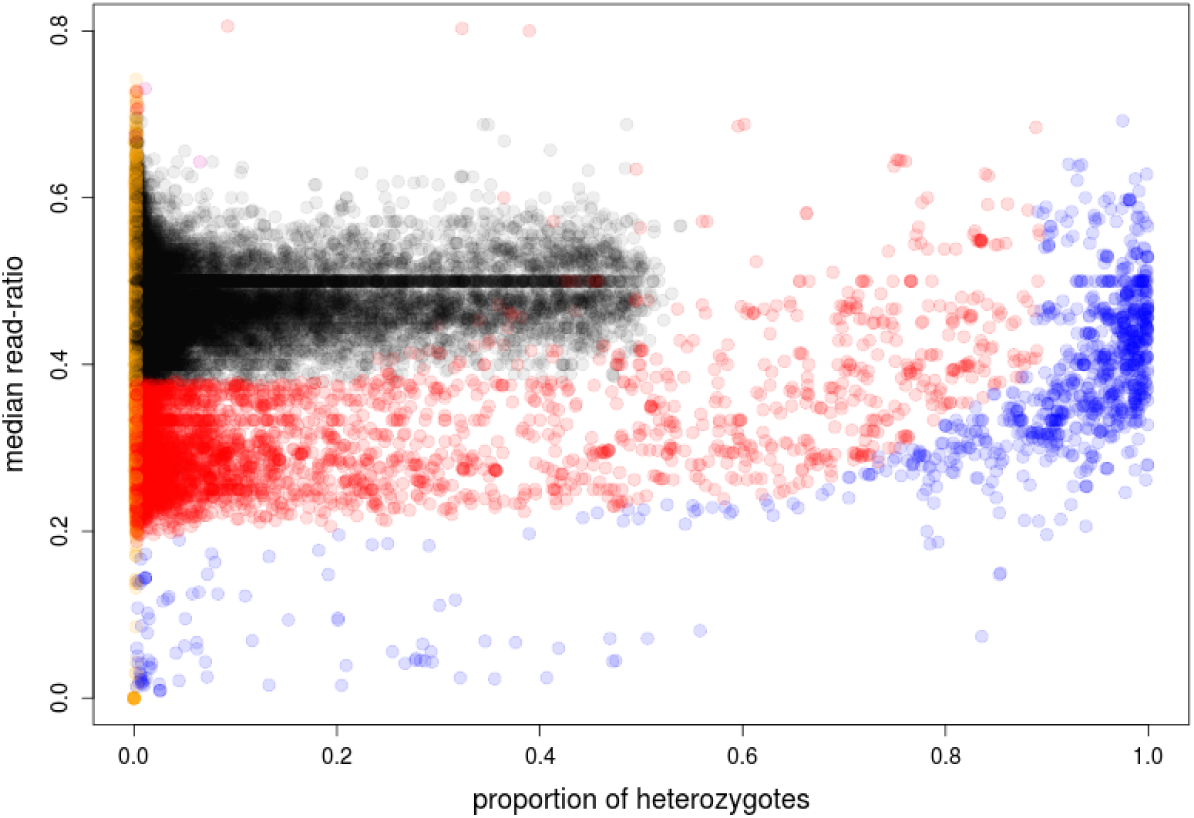
Characterization of duplication effect over the SNP dataset. The bivariate scatter plot display of the distribution of the 26,005 SNPs with the median read-ratio deviation of heterozygotes (y-axis) plotted against the proportion of heterozygotes (x-axis). The median read-ratio describes the deviation from equal alleles read-ratio (50/50) expected for heterozygotes. Black, red, blue and orange points represent singletons, duplicated, diverged and low confidence SNPs, respectively. Here, we only represent the most informative graphical representation of SNPs classification, which is derived from the graphical pattern of SNP categories (i.e. singleton, duplicated, diverged) demonstrated by McKinney et al. (2017a) with data simulations as well as empirical analyses, Other graphical representations are presented in figure S1.

### 3.2 Measuring genetic differentiation

Pairwise genetic differentiation among the 21 sampling sites based on the 13,854 SNPs was extremely weak with *F*_*ST*_ values ranging from 0 to 0.00086 (average pairwise F_ST_ = 0.0001 with 93% non-significant pairwise F_ST_ values). In contrast to SNPs, the genetic differentiation estimated using the *V*_ST_ index across the 1,521 CNV loci exhibited a stronger and significant signal of genetic differentiation, with a value ranging from 0.01 to 0.10 (average pairwise *V*_ST_ = 0.04 and all pairwise values were significant).

PCA based on individuals’ SNP genotypes did not reveal any significant pattern of genetic differentiation among the 21 sampling sites (figure 1b). In contrast, PCA based on individuals’ CNV data revealed a strong spatial structure that does not correspond to the geographic proximity among sampling sites (figure 1c). Based on the first principal component, which explained 5.31% of the variance, we observed that the majority of lobsters sampled from sites at both edges of the sampling area (i.e. individuals originated from sampling sites labelled as T1P., T5P. and two of T2P., which are displayed by color codes in blues, purples and dark green respectively) were more similar to each other than to individuals collected from central sites (T3P. and T4P. sites). Furthermore, this spatial discontinuity did not reflect the shape and direction of the main current pattern observed in this region and illustrated in the figure 1a.

### 3.3 Genotype - environmental association

Strong correlations were observed among the five thermal parameters initially considered (figure S2). Hence, we only selected two SST variables (*i*.*e*., SST annual minimum and annual variance) as predictors for the RDA analysis (SST values detailed in table S1). First, we selected the SST annual minimum because this predictor showed the lowest level of collinearity among the five thermal parameters considered. Moreover, this parameter has been associated with thermal adaptation in the American lobster at a larger geographical scale than studied here (Benestan et al., 2016). Second, because most marine species are adapted to less thermal variability compared to terrestrial environments (Sunday et al., 2011), we considered that the SST annual variance may be the factor that most affects fitness-related traits (e.g. growth and survival) of the American lobster. In particular, the SST variance parameter reflects the degree of unpredictability of thermal conditions (e.g. extreme thermal events) that pelagic larvae may face. Furthermore, Larouche & Galbraith (2016) also observed that the southern part of the Gulf of St. Lawrence experiences the highest thermal amplitude of the Canadian marine coastal waters.

No association between environmental variables and SNP genetic variation was detected by the RDA (i.e. ordistep adj. R^2^ = 0 for both SST annual minimum and SST annual variance) and the LFMM 2 analyses (data not shown). In contrast to SNPs, both SST annual variance and SST annual minimum were significantly associated with CNV variation across sampling sites based on the forward selection procedure (i.e. *ordistep* function). We also observed a stronger association for the SST annual variance (adj. R^2^ = 0.105; *P* < 0.001) than the for SST annual minimum (adj. R^2^ = 0.066; *P* < 0.001). The RDA conducted with selected predictor variables (i.e. SST annual variance and SST annual minimum) and the CNV dataset was highly significant (*P* < 0.001; ANOVA 1,000 permutations) and explained 2.4% of the total CNV variation (adj.R^2^ = 0.024). A total of 83 loci were detected as candidate outliers, where 67 and 16 were associated with SST annual variance and SST annual minimum, respectively. No CNV candidates were associated with both SST variables.

Linear mixed-effects models detected 288 CNV loci significantly associated with annual SST variance (adj. *P* < 0.01). Among these 288 CNVs, 48 were also identified from the RDA (i.e. 15.6% in common among all outliers detected with both methods combined). Although the LME models returned 19 CNV loci significantly associated with the SST annual minimum, none were in common with the 16 candidates identified by the RDA. *V*_ST_ estimation for all 1,521 CNV loci revealed nine outlier loci with substantially elevated levels of population differentiation (i.e. *V*_*ST*_ values above a threshold of six standard deviations), and all of them were identified in association with the SST annual variance.

The PCA performed with the set of 48 CNV candidates associated with the SST annual variance showed that sampling locations formed two distinct groups that were mainly discriminated by the first principal component (PC1), which explained 50.20% of the variance (figure 3a). Sampling sites located at both edges of the sampling area (i.e. T1P1, T1P2, T1P3, T1P4, T2P1, and T2P2 in the north; T5P1, T5P2, T5P3 and T5P4 in the south) were clearly separated from those sampling locations from the center part of the studied range (i.e. T2P4, T2P5, T3P1, T3P2, T3P3, T3P4, T4P1, T4P2, T4P3 and T4P4). The sampling site T2P3 displayed an intermediate position. Based on a Broken-stick model, only the first principal component was considered for interpretation. The geographical transposition of PC1 scores obtained from sampling sites centroids within the climate layer of the annual SST variance showed a strong association between the thermal environment and CNV variation (figure 3b). The cluster represented by sites from the center-range of the sampling area corresponded to geographical areas with high variance of SST, while the second cluster contained sampling sites from the edges of the sampling area with low variance of SST. Although the Broken-stick model supported only the first PC, we also observed that PC2 separated sites located at both edges of the sampling area, albeit to a lesser degree than the separation observed on the first axis (figure 3a).

**Figure 3:**
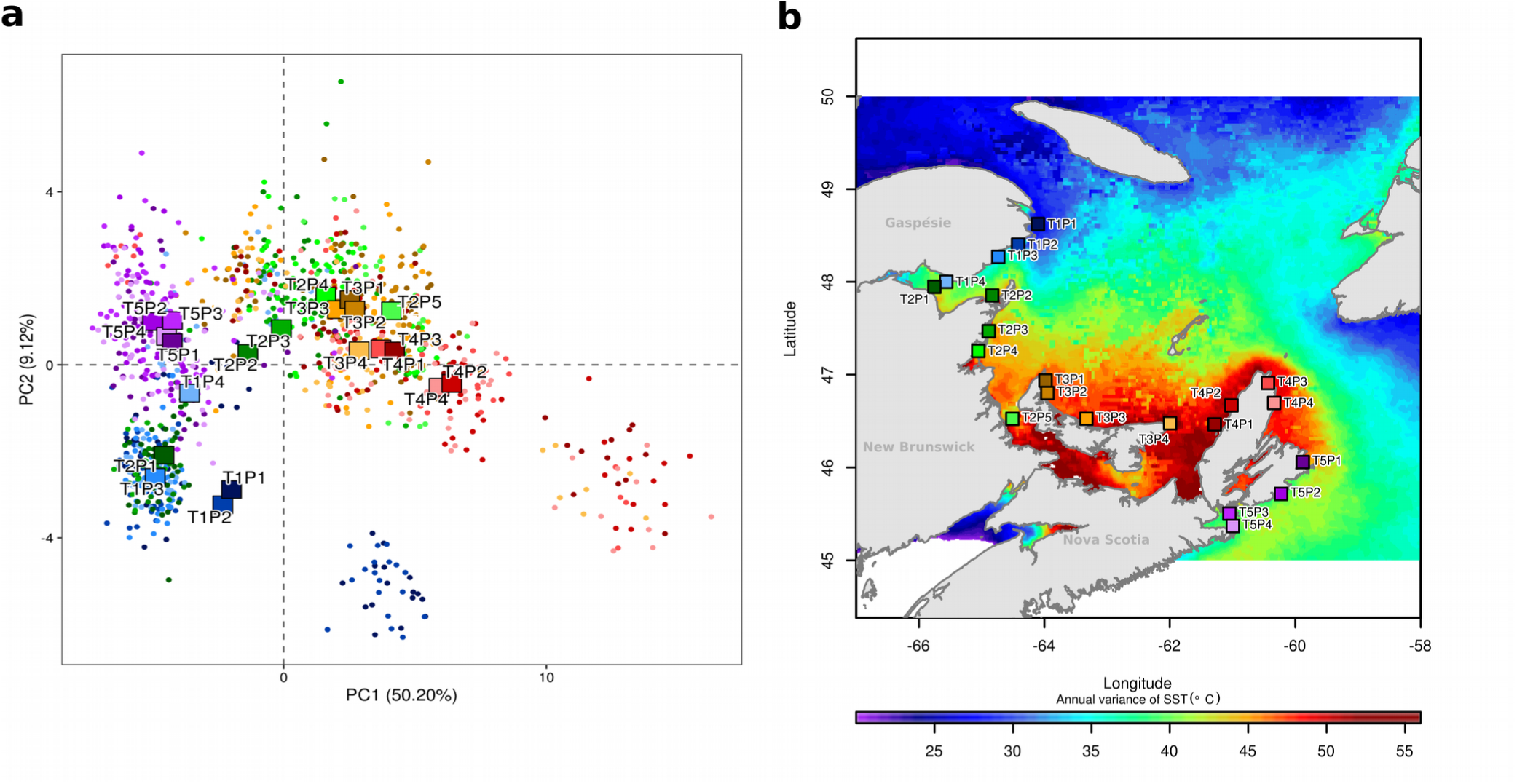
PCA based analysis of the environmental associated CNVs. PCA was applied to the normalized read depth matrix of 48 CNV loci associated with SST annual variance. **(a)** PCA biplot showing individuals (points) and sampling sites (squares) positioned using the centroid value. Note the clustering of sampling locations based on PC2 (namely those labelled as T1P. and T5P.) despite the geographic separation). Note also the subclustering of the two sampling sites T1P1 and T1P2, which are also visible though modal clustering of CNVs profiles displayed in the figure 4. **(b)** Climate map depicting the annual variance of sea surface temperature within the south of the Gulf of St. Lawrence, Canada. Sampling sites are represented by squares. For both PCA biplot and climate map, individuals and sampling sites are colored according matching the color scale used in the Figure 1.

The shape of the relationship between read depth (normalized) at all 48 CNV candidates and SST annual variance was relatively similar for the majority of the 48 candidate CNVs (figure 4 and figure S4). Indeed, analyses based linear mixed-effect models revealed that this relationship was non-linear for 43 loci and was best described by a Gompterz function or a logistic function for 35 (73%) and 8 (16%) CNV loci respectively (figure S5). Furthermore, a correlation analysis of read depth among the 48 CNV outlier loci suggested that a block of these CNVs could possibly be physically linked whereas lower correlation was observed among other CNVs(figure S6). Thus, the correlation (R^2^) among 35 out of the 48 candidate CNVs was generally over 0.40 (median R^2^ = 0.42; R^2^ ranging from 0.12 to 0.84) whereas it was generally less than 0.10 for the others. In absence of a lobster genome to map our sequence reads, this pattern suggests that most of these candidate CNVs are possibly located within a same chromosomal region whereas the remaining could be distributed on other chromosomes. For most of the 43 CNV loci best described by a non-linear function, read depth remained stable until a SST annual variance value of ∼45 °C^2^, and then drastically increased and stabilized again at annual variance value of ∼50 °C^2^ (figures 4 and S4).

**Figure 4:**
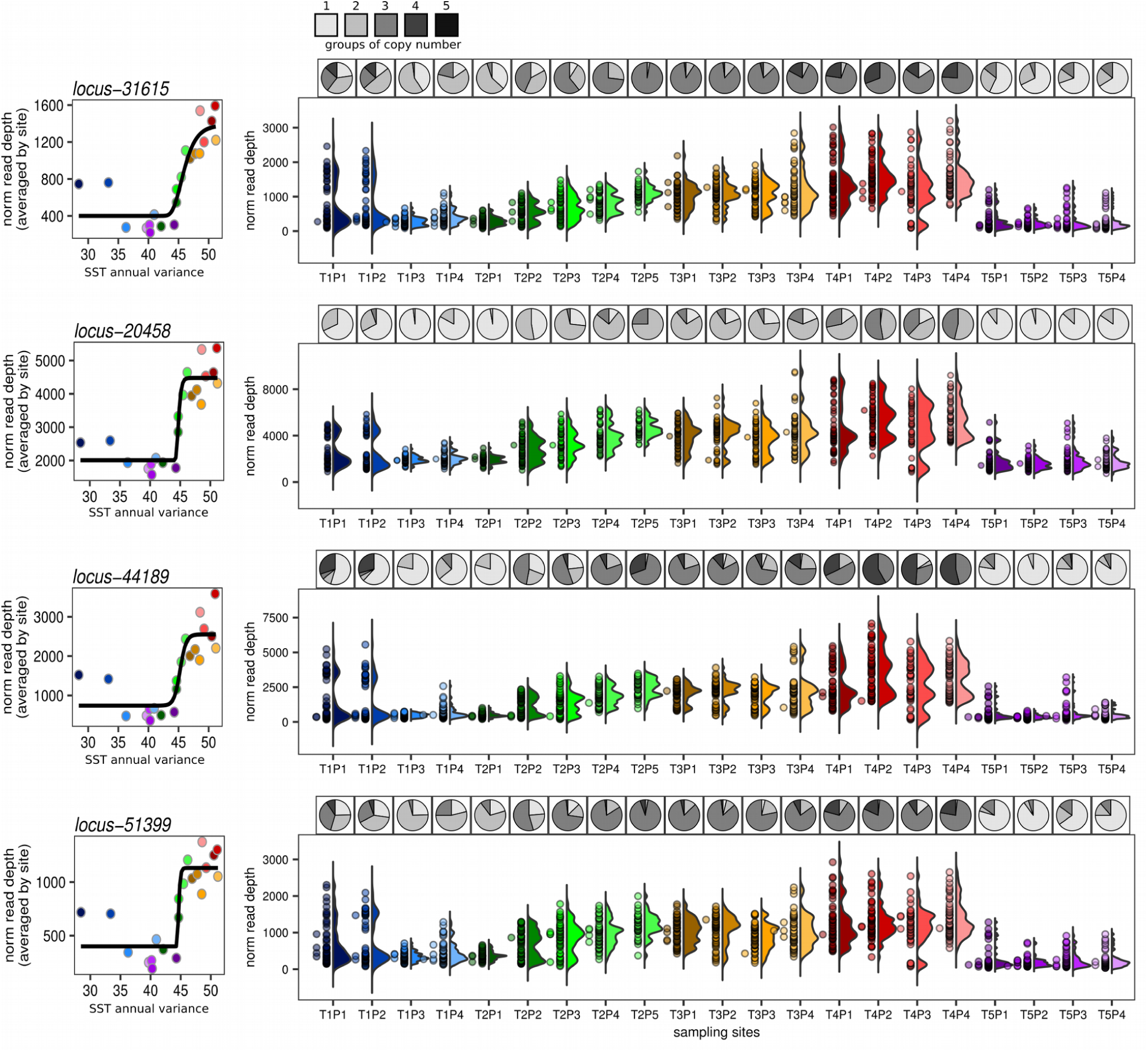
Overview of CNV profiles for the four most highly differentiated loci (*V*_*ST*_) associated with the SST annual variance. (Left column panel) Correlation plot between SST annual variance and the average read depth observed at each sampling site. Black lines represent the Gompertz fit model for each locus. (Right column panel) For each sampling site (x-axis), half-violin plots represent the density of normalized read depth distribution (y-axis) while points represent individual lobsters. The colour for the different sites is according to their respective color code depicted in the figure 1a. Header pie charts represent the proportion of individuals classified within each putative ‘copy group’ (from 1 to 5) among each sampling site. For each CNV locus the putative number of “copy groups” was estimated based on independent EM algorithm (see M&M for details). White to black box-scale represents the different groups of copy number.

Based on the results above, we defined two main clusters: (i) the first consisting of sampling sites experiencing low values of SST annual variance (i.e. σ^2^_Temp_ < 45°C^2^), and (ii) the second one consisting of sampling sites experiencing high values of SST annual variance (i.e. σ^2^_Temp_ > 50 °C^2^). The level of genetic differentiation estimated using the 1,521 CNV loci was higher between sampling sites originated from the two distinct clusters (i.e. edges cluster vs. central cluster) than sampling sites from the same cluster. The average of pairwise *V*_ST_ values were 0.0548 (SD = 0.0131) and 0.0298 (SD = 0.0149) for *between clusters* versus *within cluster* respectively, and the Kolmogorov-Smirnov test showed that this difference was significant (KS: D = 0.616, P < 0.001).

### 3.4 Discrete groups of CNVs

The identification of discrete groups of copy number categories was examined using the model-based clustering procedure for each of the 48 CNVs loci that was associated with the annual variance of SST. This analysis revealed that among these 48 candidate CNVs, up to five discrete categories can be delineated, where each category may represent a specific number of copies for a given locus (see figure S7, S8 and S9 for details). While the first copy number group (the one best supporting a unimodal distribution of coverage) could possibly represent individuals with a single (presumably diploid) copy for a given locus, we prefer to remain conservative in absence of a reference genome to definitively confirm this. We therefore prefer referring to copy number group 1 to 5, with 1 representing the group with the lowest copy number and 5, the highest. We observed a highly significant difference in the distribution of copy number categories among sampling locations(Wilcoxon test, P < 0.001). Generally speaking, for each of these 48 candidate CNVs, low-level copy number categories were mainly found in sampling sites characterized by low SST annual variance, which are located at the two edges of the sampling study area. By contrast, groups with higher level of copy numbers were located in sampling sites with high SST annual variance within the central part of the study area (figures 4 and S8).

## Discussion

Our study reveals that CNVs represent a non-negligible fraction of genetic variability in the American lobster genome, and that this variability can be detected with reduced representation sequencing data using McKinney’s et al. (2017a) framework. The level of genetic differentiation among the 21 sampling sites was significantly and markedly higher with CNVs than with SNPs. More importantly, we did not detect associations between SNPs and any of the temperature variables investigated, but did find a strong and significant pattern of association between a set of CNVs and sea surface temperature variance, which did not reflect geographic proximity. This suggests that variation in copy numbers at those loci could be involved in local (thermal) adaptation. Overall, our study supports the view that CNVs represent a significant portion of genetic polymorphism, and in turn, are relevant genomic markers for population genomic studies.

### On the importance of detecting CNVs

Reduced-representation shotgun sequencing (RRS, which include RADseq, ddRAD, GBS and Rapture) is a powerful tool for non-model systems and large-scale studies, since it allows sequencing a large number of DNA sequences for thousands of samples simultaneously at a minimal cost (Davey et al., 2011). However, loci present in multiple copies are - even if the number of copies does not vary between individuals - often confounding for DNA sequence alignment methods and can affect polymorphism inferences, leading, for instance, to errors in SNPs calling and bias in allele frequency estimation (McKinney et al., 2017a). In this study, we improved confidence in SNP calling from RRS (Rapture) sequencing data by classifying and considering separately loci with copy-number variability, the CNVs, based on the methodology developed by McKinney et al (2017a) (i.e. HDplot). Beyond providing a more polished SNP dataset, our classification allowed us to analyze a new dimension of genetic variability, a CNV dataset with the same original RRS sequencing data.

Simulations and empirical data by McKinney et al. (2017a) support the robustness of this approach, based on a set of simple summary statistics of SNP data, by showing that it correctly identified duplicated sequences with >95% concordance with loci of known copy status. While the stochastic nature of RRS data, especially at low to moderate sequencing depth, can affect the estimations of the read ratio deviation (i.e. the main statistic of the HDplot approach; McKinney et al., 2017a), this should be minimal in our study since the Rapture approach enabled us to sequence 384 RRS captured libraries on the same sequencing chip, reducing unbalanced representation of individuals by a high rate of multiplexing. Moreover, genotype calling quality was assured by a high average read-depth per locus (average across all singletons SNPs was ∼28X reads/locus), leading to very few missing data over individual genotypes. The interpretation of variation in CNV profiles among study sites from the distribution of normalized read depths was also improved through the delineation of different discrete groups of copy number using a model-based clustering approach. This approach, which includes the *Mclust* procedure used here (Fraley & Raftery, 2012) or other methods such as Bayesian mixture model-based clustering (Medvedic et al., 2004), offers advantages over heuristic methods (e.g. k-means clustering), such as the ability to measure uncertainty about the resulting clusters and to formally estimate the number of clusters (Fraley & Raftery 2007). Furthermore, model-based clustering procedures have been broadly used in genetics and ecology. For instance in studies using microarray to document patterns of gene expression or cancer cell type identification, as well as life-history traits variation (Freyhult et al., 2010; Côté et al. 2015; Barajas-Olmos et al., 2019). Altogether, the set of methods presented in this study thus provides a way to maximize the genetic information drawn from RRS sequencing by improving the accuracy of the SNP dataset and by providing a CNV dataset. Although the approach faces the same limits as other CNV detection methods based on SNPs arrays or whole-genome sequencing (e.g. mappability issues, GC content, PCR duplicates, DNA libraries quality; Teo et al., 2012), it also provides the means to consider CNVs in non-model species even without extensive genomic or financial resources. Here, for instance, with a genome size of about 4.5Gb (Jimenez et al., 2010), and in absence of a reference genome, it would have essentially been impossible to screen for CNV variation across the entire genome for hundreds of individuals.

### Copy number variants represent a substantial fraction of genetic diversity in the lobster genome

By applying the above-described classification methods, we identified that ∼20% of the loci successfully sequenced represented copy number variants in the American lobster. Variation in copy-number resulting from whole genome duplication, transposable elements, or duplication, deletion and rearrangements of various repeated DNA has been implicated in the change of genome size among eukaryotes (Biémont, 2008; Cioffi & Bertollo., 2012; López-Flores & Garrido-Ramos, 2012). For example, Prokopowich, Gregory and Crease (2003) showed a strong relationship between rDNA copy number and the genome size of 162 species of plants and animals. Studies of genome evolution among taxonomic groups represented by species exhibiting some of the highest genome sizes in eukaryotes such as Coniferae (Nystedt et al., 2013) and Amphibians (Sun & Mueller, 2014), revealed that the proliferation of repetitive elements represent the major mechanism driving large genome size. For instance, it has been estimated that up to 40% of the Mexican axolotl *(Ambystoma mexicanus*) 32 Gb genome is represented by repetitive DNA sequences, and that in the tiger salamander (*Ambystoma tigrinum*) this proportion may constitute up to 70% of the genome (Keinath et al., 2015). Thus, it is plausible that the large (∼4.5 Gb) genome of American lobster is also made up of an important proportion of repeated regions, including the CNVs detected in this study. This is supported by recent work on other crustacean genomes, which revealed prominent proportions of repeat-rich regions (Song et al., 2016, Verbruggen 2016; Yuan et al., 2018, Fang et al., 2019). For instance, up to 23% of the genome (∼1.66 Gb) of the Pacific white shrimp, *Litopenaeus vannamei*, has been reported as comprising repeated DNA sequences (Zhang et al. 2018). Our correlation analysis of read depth among the 48 CNV outlier loci suggested that 35 out of the 48 candidate could possibly located within a same chromosomal region (given that R^2^ varied between 0.12 and 0.84 between them) whereas the remaining could be distributed on other chromosomes (R^2^ generally less than 0.10). Admittedly however, a firm confirmation of this pattern must await the production of a high quality reference genome for the American lobster. Nevertheless, the fact that we identified that ∼20% of the loci successfully sequenced represented copy number variants and that their geographic pattern of variation is associated with that of annual temperature supports the view that CNVs represent a significant proportion of the variation present in the American lobster’ genome and raises the hypothesis that this variation could play important a role in local adaptation, as discussed below.

### CNVs reveal a stronger genetic structure than SNPs

Congruent with previous population genomic data for lobsters from the southern Gulf of St. Lawrence (Benestan et al 2015), we observed an extremely weak level of genetic structure at SNP markers. Our results are also consistent with the weak genetic structure (F_ST_ typically less than 0.01) generally observed using SNPs in other marine organisms (fishes, DiBattista et al., 2017; Junge et al., 2019; molluscs, Sandoval-Castillo et al., 2018; crustaceans, Al-Breiki et al., 2018; echinoderms, Xuereb et al. 2018). This weak differentiation likely reflects the combination of pronounced gene flow and large effective population size among lobsters from the 21 sampling sites. Bio-physical larval dispersal models support high levels connectivity between lobster populations caused by high dispersal of lobsters during early life stages (dispersal distances ranging from 5-400 km; Chassé & Miller 2010; Quinn et al., 2017). In addition, a recent mark-recapture study revealed that benthic movements at the adult stage also likely contribute to high gene flow and high connectivity for this species (Morse et al., 2018).

In contrast to SNPs, CNVs showed markedly higher levels of differentiation in the study area (*i*.*e*. average *V*_*ST*_ = 0.043 and all pairwise *V*_*ST*_ were significant). Although a direct comparison between *F*_*ST*_ and *V*_*ST*_ values cannot be made as the two metrics rely on very different data, our study nevertheless suggests that CNVs unveil population structure that was not detected using SNPs. This difference in structure is consistent with previous studies in humans showing that CNVs contribute to higher genetic divergence than SNPs (Levy et al., 2007; Pang et al., 2010; Sudmant et al., 2015). Furthermore, recent studies in marine fishes also showed that genomic regions including structural variants account for higher genetic divergence than SNPs that are distributed across the whole genome (Berg et al., 2016; Barth et al 2019; Kess et al., 2020, Cayuela et al., 2020). Several hypotheses may explain this pattern. For instance, SVs such as inversions or some categories of CNVs locally limit recombination (Rowan et al., 2019), thus reducing genome homogenization despite gene flow. Some CNVs, such as transposable elements, exhibit high evolutionary dynamics and have important implications for rapid genome evolution and adaptation (Kofler, Nolte and Schlötterer, 2015; Rey et al., 2016). By affecting a larger fraction of the genome, or by having stronger effects on the phenotype, CNVs may be particularly sensitive to positive or negative natural selection. Although the mechanisms remain poorly known the increasing evidence that CNVs sometimes exhibit more differentiation than SNPs leads us to consider CNVs as powerful genetic markers for characterizing spatial genetic structure and to take advantage of present technology to include them in future population genomic studies. Finally, we can ask whether CNVs recapitulate more structure or more diversity simply because they have wider amplitude of variation than bi-allelic SNPs or whether it is because they capture a fundamentally different aspect of genomic variation. Other techniques have also recently been proposed to better leverage RRS data to improve population discrimination power by combining multiple SNPs per RAD locus (e.g. microhaplotypes; McKinney et al. 2017b). In the future it will be interesting to compare the relative strengths of CNV and microhapotype markers for detecting structure and whether or not they recover similar or different patterns of population structure.

### CNV-environment associations are stronger than SNP-environment associations

Although redundancy analysis (RDA) and LFMM are efficient methods to identify candidate SNPs associated with variability in environmental conditions (Forester et al., 2018; Caye et al., 2019), no significant relationship was detected between any of the SNPs and the temperature variables tested. This result differs from the findings of Benestan et al. (2016), who reported a strong association between 505 SNPs and thermal conditions (i.e. minimum annual SST) in the American lobster. However, the geographic scope of their study covered most of the species’ distribution in the Northwestern Atlantic, and like several other Northwestern Atlantic species (i.e. Atlantic cod - *Gadus morhua*, European green crab - *Carcinus maenas*, Northern shrimp - *Pandalus borealis*, and the Sea scallop - *Placopecten magellanicus*) (Stanley et al., 2018), the genotype-environment associations were predominantly driven by a north vs south dichotomy (Benestan et al. 2016), while the southern part of the study area of those studies was not covered here.

Our analyses identified 48 CNVs (out of 1,521), significantly associated with the SST annual variance, supporting the hypothesis that variation in copy numbers at those markers could be associated with local adaptation to temperature variance in the American lobster. The role of CNVs in local adaptation has been recognized and understood in plants, animals, and fungi (e.g. Gonzalez & Petrov 2006; Bazzicalupo et al., 2019; Nelson et al., 2018; Prunier et al., 2019; Zhang et al., 2020). In marine fishes, CNVs have been associated with freeze resistance (Hew et al., 1988; Hayes et al, 1991, Chen et al., 2008; Desjardins et al., 2012). For instance, Hayes et al. (1991) demonstrated antifreeze proteins (AFPs) gene copy number and arrangement among certain populations of winter flounder (*Pseudopleuronectes americanus*) along the Northeastern Atlantic coast. Similarly, Desjardins et al. (2012) found that multiple copies of AFP genes improved gene-dosage effect and transcription level between two closely related wolffish species (*Anarhichas lupus* and *A. minor*), facilitating the colonization of a shallow-water habitat where the risk of freezing is elevated. Finally, Martinez Barrio et al., (2016) demonstrated the implication of copy number variants in local adaptation to varied salinity habitats in Atlantic Herring (*Clupea harengus*) using a combination of genomics approaches (i.e. genome assembly, pool sequencing and SNP chip analysis). By showing empirical evidence for CNV-environment association, our results thus bring additional evidence for the importance of structural variation for local adaptation in marine species.

The repeated observation of a similar relationship among 48 CNV loci (similar shape of the regression model) associated with the annual variance of SST raises questions about the nature of this mechanism within the genome. As explained above, our results suggest that the majority of CNV loci associated with the SST annual variance could be physically linked, perhaps within a same chromosomal region. However, the inherent features of our sequencing approach, which represents only a reduced sample of the entire genome, coupled with the absence of a reference genome for American lobster, does not allow us to rigorously address this issue and refute alternative explanations, such as a strong effect of selection driving to covariation in copy numbers among different CNVs. Future works should be able to address this by taking advantage of technological advances in long-read sequencing (i.e. Nanopore or PacBio technologies) as well as the decreasing costs to develop references genomes in non-model species.

In contrast with some congruence observed between the sets of CNVs identified by the two GEA approaches, we observed a complete lack of overlap between results obtained for the association with annual minimum SST. We can only speculate on the possible causes for this. First, the initial forward-selection model (RDA) showed that the association strength (i.e. given by the R^2^ value) of the SST annual minimum was twice lower than the SST annual variance, which is corroborated by the small number of CNV candidates detected by the RDA in association with the SST annual minimum. Second, LME models only identified a small number of CNV candidates associated with the SST annual minimum. Overall, this lack of overlap may suggest that this environmental variable could be inappropriate to examine adaptive genetic variation at the spatial scale of our study. Alternatively, it is not unusual to observe limited overlap of markers identified as potentially under selection among different genome scan or GEA methods (Gagnaire et al. 2015).

The details of how CNVs may underlie locally adaptive traits implicates various complex genomic mechanisms, such as the remodeling chromatin architecture (Dunaway et al., 2016) and the modulation of gene expression (i.e. gene-dosage effect; Desjardin et al., 2012, Zhang et al. 2019). The alteration of gene-dosage balance may negatively impact individual fitness, this has been mostly documented in relation to diseases (Gamazon & Stranger 2015; Rice & McLysaght., 2017). However, other studies showed that certain gene-dosage effects caused by CNVs can be positive and facilitate the adaptation of organisms to environmental changes. For example, CNVs are thought to drive insecticide resistance in populations of *Aedes* mosquitoes by increasing the effectiveness of detoxification enzymes (Faucon et al., 2017, Weetman et al., 2018). Furthermore, multiple gene copies are important drivers of differences in gene expression between populations of three-spined stickleback (*Gasterosteus aculeatus*), which cause phenotypic variation that affects habitat-specific selection to contrasted environments (Huang et al., 2019).

Interestingly, we found that it was the variance of environmental conditions (i.e. SST annual variance) that was best associated with CNVs instead of the more commonly investigated average, minimum, or maximum temperature values. It is well documented that many organisms often use phenotypic plasticity as a strategy for maximizing their fitness in variable environments (Schlichting & Pigliucci 1998; Aubin-Horth & Renn, 2009; Laporte et al., 2016). Extremes in variation of annual temperature are stressful for organisms and increased gene-copy number may provide the capacity for adjusting gene expression to survive in the face of these extreme events. Thus, one would predict that an adaptively plastic genomic mechanism would imply populations that experience highly variable temperatures evolve toward increased copy number of relevant genes. Given the close interplay of transposable elements (a common type of CNV) and DNA methylation silencing of their expression (Kelleher et al., 2020), the expression of such genes could be controlled by environmentally-mediated changes in DNA methylation. Indeed, DNA methylation is extremely labile and greatly affects gene expression (McCue et al., 2012; Rougeux et al., 2019). For instance, DNA methylation can change in minutes following an environmental change and return in their initial configuration in days in previous environmental conditions (Huang et al., 2017). Furthermore, environmental stress can also induce a reactivation of transposable elements via DNA demethylation, which could relatively quickly result in an increase of CNVs through the genome that will be submitted to selective pressures in the new environment (Rey et al., 2019). Admittedly, the possible mechanisms by which variation in CNVs in association with thermal variance could indeed be involved in local adaptation remain hypothetical here. Nevertheless, our observations argue strongly for the value of investigating such mechanisms in future studies.

### Conclusion

Our study highlights the importance of structural variants such as CNVs in molecular ecology by revealing their implication in shaping population structure and local adaptation in a marine species. Moreover, our contribution provides an approach to characterize and analyze CNVs with reduced-representation sequencing, an approach which is applicable with limited financial resources and without reference genome. We believe that despite some limitations, which are identified above, this approach will enable researchers to take advantage of CNV markers to better characterize population structure of non-model species for conservation or management. Moreover, extending CNV characterization and analysis to a wider range of species and ecological contexts will allow better understanding of the ecological and evolutionary significance of these structural variants.

## Data accessibility

(1) The following datasets generated for this study and available from the Dryad Digital Repository: doi:10.5061/dryad.vt4b8gtnv

- VCF file of no filtered and filtered SNPs dataset
- Matrix of normalized read depth for CNV loci
- Estimated statistics for SNPs duplicated effect detection
- Series of R scripts (environment data reduction, CNVs identification, read-depth normalization)
- The demultiplexed DNA sequences in FASTQ format have been archived on NCBI at : ###

## Authors’ contribution

L.B and R.R designed and supervised the project. Y.D conducted literature mining and laboratory work. Bioinformatics and data analyses were conducted by Y.D with input from E.N., M.L. H.C. K.W. and Q.R. Original draft preparation was led by Y.D. and all authors contributed to the writing and editing of the final version of the manuscript.

## Competing interest

None declared

## Acknowledgements

We thank scientists from the Department of Fisheries and Oceans and Canadian fishers who helped collecting the samples. We are grateful to Alison Devault and the Arbor Biosciences team for DNA probes synthesis and methodological advices. We also thank the personnel of the IBIS sequencing platform for their assistance in developing the Rapture assay for highly multiplexed configuration. We thank three anonymous reviewers and the associate editor for their insightful reviews and very constructive comments and suggestions. This research was financially supported by a Strategic Partnership Grants for Projects from the Natural Sciences and Engineering Research Council of Canada to LB and RR (Grant number STPGP 462984-14).

## Supporting information

Additional supporting information may be found in the online version of this article

Table S1: Geographic coordinates, sample size (i.e. number of samples successfully genotyped), sequencing effort, environmental parameters across the 21 sampling locations.

Table S2: Filtering sequencing data - setup pipeline

Table S3: SNPs classification results

Figure S1: Characterization of duplication effect over SNP dataset

Figure S2: Correlation plots of temperature parameters

Figure S3: Impact of read depth normalization for CNVs data

Figure S4: Relationship between SST annual variance and CNVs read depth

Figure S5: Model comparisons

Figure S6: Heatmap of statistical relationship among the 48 CNV loci associated with the SST annual variance

Figure S7: Discretization of copy groups for 48 CNVs associated with SST annual variance

Figure S8: Mclust model performance among the 48 CNVs locus associated with SST annual variance.

Figure S9: Mclust model features for the 48 CNV loci associated with SST annual variance.

## Notes

### Competing Interest Statement

The authors have declared no competing interest.

