## Supplementary Material for "Copy number variants outperform SNPs to reveal genotype-temperature association in a marine species"

Table S1 : Geographic coordinates, sample size (i.e. number of samples succesfully genotyped), sequencing effort, environmental parameters across the 21 sampling locations.

| Sampling Site | Latitude | Longitude | Sample size | Sequencing effort* | | Sea Surface Temperature (annual) | | | | |
|---|---|---|---|---|---|---|---|---|---|---|
| | | | | Median | sd | minimum | maximum | variance | mean | range |
| T1P1 | 48.6217 | -64.1036 | 53 | 391,723 | 109,456 | -0.5 | 14.7 | 28.4 | 5.8 | 15.2 |
| T1P2 | 48.4000 | -64.4193 | 55 | 431,657 | 120,108 | -0.1 | 15.9 | 33.4 | 6.4 | 16.0 |
| T1P3 | 48.2687 | -64.7353 | 50 | 390,123 | 115,093 | -0.2 | 16.4 | 36.3 | 6.7 | 16.5 |
| T1P4 | 48.0001 | -65.5648 | 53 | 406.794 | 145,077 | -0.8 | 16.8 | 41.0 | 7.1 | 17.6 |
| T2P1 | 47.8545 | -65.7533 | 39 | 372,618 | 99,541 | -0.6 | 17.0 | 42.1 | 7.2 | 17.5 |
| T2P2 | 47.8125 | -64.8319 | 55 | 398,611 | 94,948 | -0.6 | 17.7 | 44.7 | 7.4 | 18.3 |
| T2P3 | 47.4677 | -64.8869 | 56 | 432,681 | 138,011 | -1.1 | 17.5 | 44.7 | 7.6 | 18.6 |
| T2P4 | 47.2553 | -65.0533 | 56 | 471,803 | 123,672 | -0.9 | 17.7 | 45.5 | 7.8 | 18.6 |
| T2P5 | 46.5230 | -64.5100 | 36 | 433,754 | 109,977 | -0.9 | 17.6 | 46.2 | 7.8 | 18.5 |
| T3P1 | 46.9363 | -63.9825 | 55 | 564,877 | 219,624 | -0.8 | 18.2 | 46.9 | 8.0 | 19.0 |
| T3P2 | 46.8019 | -63.9519 | 52 | 489,805 | 181,699 | -0.6 | 18.4 | 47.8 | 8.0 | 19.0 |
| T3P3 | 46.5238 | -63.3311 | 55 | 438,196 | 151,902 | -0.7 | 18.7 | 48.5 | 8.0 | 19.3 |
| T3P4 | 46.4658 | -61.9952 | 53 | 393,938 | 129,041 | -1.0 | 19.0 | 51.2 | 7.5 | 19.9 |
| T4P1 | 46.4638 | -61.2844 | 53 | 472,244 | 187,437 | -0.4 | 19.1 | 50.5 | 7.7 | 19.4 |
| T4P2 | 46.6685 | -61.0167 | 55 | 459,954 | 157,628 | -0.8 | 19.1 | 51.1 | 7.6 | 19.9 |
| T4P3 | 46.9095 | -60.4350 | 45 | 451,232 | 173,327 | -0.9 | 18.7 | 49.2 | 7.0 | 19.6 |
| T4P4 | 46.6952 | -60.3368 | 54 | 364,437 | 86,385 | -1.0 | 18.7 | 48.6 | 7.0 | 19.7 |
| T5P1 | 46.0581 | -59.8795 | 49 | 472,664 | 144,766 | -0.4 | 18.3 | 44.3 | 6.8 | 16.7 |
| T5P2 | 45.7156 | -60.2260 | 52 | 409,104 | 190,808 | -0.4 | 17.6 | 40.3 | 6.7 | 18.0 |
| T5P3 | 45.5030 | -61.0494 | 53 | 469,551 | 279,181 | -0.5 | 17.2 | 40.2 | 7.1 | 17.7 |
| T5P4 | 45.3394 | -60.9944 | 52 | 492,587 | 231,201 | -0.7 | 17.3 | 39.7 | 7.1 | 18.0 |

*Sequencing effort corresponds to the global read count statistic (i.e. median and standard deviation of fastq files in thousands reads) summarised over all filtered individuals among each sampling site.

Table S2 : Filtering sequencing data - setup pipeline

**Sample Preparation**

- Data prepared, SNPs VCF generated and filtered using:
  - STACKS V.1.48
  - stacks workflow (last download 24 october 2019)

- Raw data cleaned with Cutadapt V.2.3.
- Samples extracted with process_radtags (part of STACKS)

**Reerence genome**

- Reference catalog of Rapture probes setup from Dorant et al. 2019
- Cleaned and demultiplexed reads aligned to ref. Catalog with:
  - bwa-mem V.0.7.17
  - samtools V.1.8
- STACKS V1.48 pipeline (with ref catalog)
  - pstacks (params: m=4)
  - cstacks (params: n=3)
  - sstacks (params: na)
  - population (params: r=0.6 ; p= 4)

**Filtering VCF output**

- STACKS VCF filtered a first time with 05_filter_vcf_fast.py (params: m=4 ; p=70 ; x=0, S=2)
- Max global missing data threshold = 15%
- Look for sample relatedness and heterozygosity problems in new VCF with vcftools
- Remove bad samples from preliminary filtering steps
- Filter this new VCF (i.e. bad samples removed) with 05_filter_vcf_fast.py (params: m=4, p=70 x=0, S=2)
- Classify SNPs into singleton, duplicated, diverged, low confidence, MAS (see supp figure S2) with
  - <u>duplicated SNPs</u>
    - Fis < -0.1 ; Fis + MedRatio < 0.1 ; Fis + MedRatio*3 < 0.78 ; Fis + MedRatio*8 < 2.3 ; MedRatio < 0.38 ; MedRatio > 0.70
  - <u>diverged SNPs</u>
    - Fis < -0.8 ; Fis + MedRatio*3 < 0.20 ; Fis + MedRatio*8 < 1.5
  - <u>low confidence SNPs</u>
    - Fis > 0.6
  - MAS (i.e. Too few samples with rare allele)
    - [N heterozygotes] + [N rare homozygotes] < 5
  - <u>singletons SNPs</u>
    - All SNPs that parse the previous filters
    - Note that all SNPs found within a potential duplicated or diverged 80bp loci were corrected as duplicated or diverged.

- Keep only SNPs that are unlinked within loci with cut-off = 0.5 (script *11_extract_unlinked_snps.py* available at https://github.com/enormandeau/stacks_workflow)

Table S3: SNPs classification results

| | Singleton | Duplicated | Low confidence | Diverged | mas |
|---|---|---|---|---|---|
| SNPs | 14,534 | 9,659 | 23 | 39 | 1,750 |
| Loci | 5,362 | 1,521 | 12 | 9 | 509 |

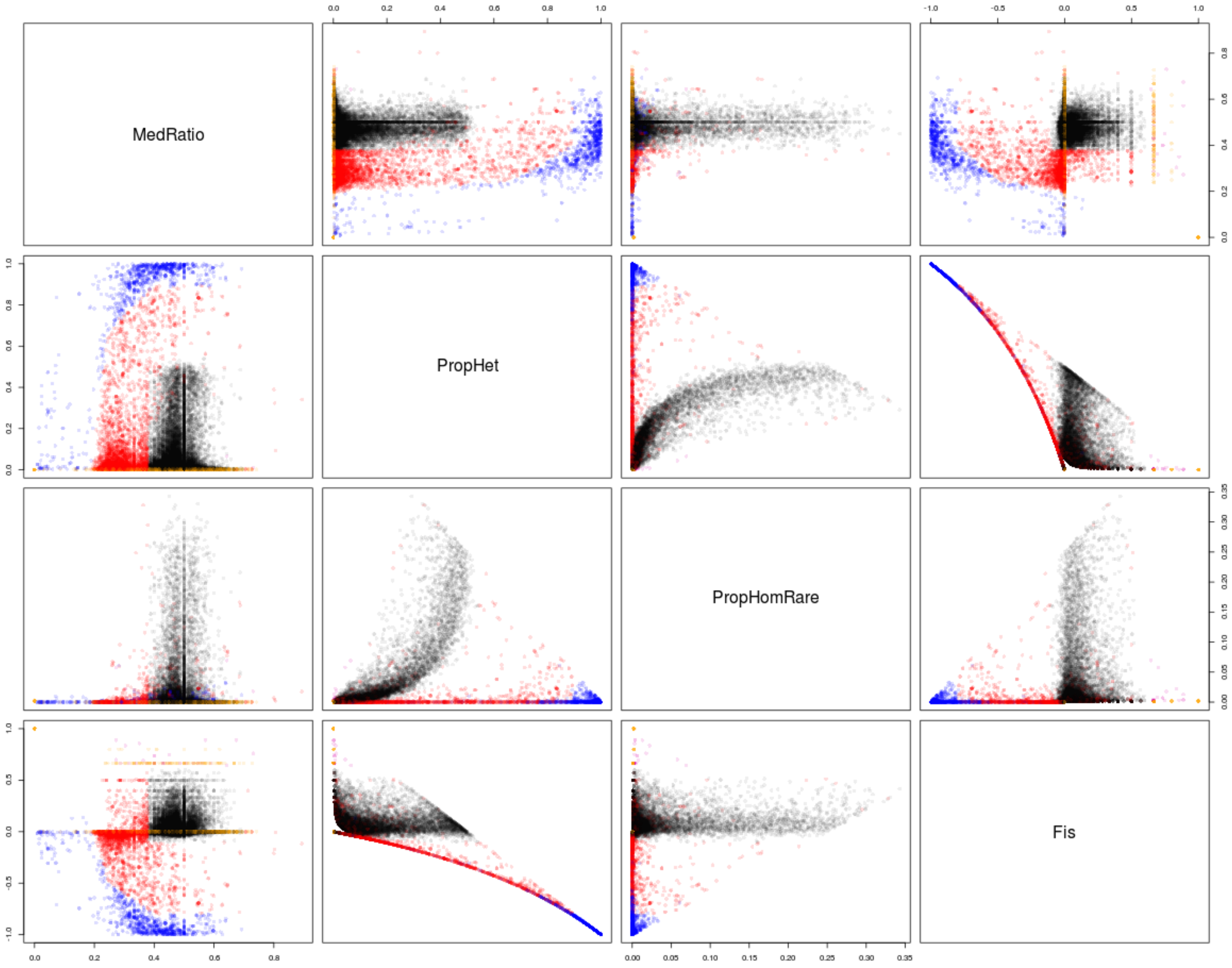


**Figure S1: Characterization of duplication effect over the SNP dataset.**
The bivariate scatter plots display the distribution of the 26,005 SNPs over four statistical parameters measured from the filtered VCF (i.e. (1) median of read allele ratio in heterozygotes (MedRatio), (2) proportion of heterozygotes (PropHet), (3) Proportion of rare homozygotes and (4) Fis). Based on the graphical patterns of SNPs categories (i.e. singleton, duplicated, diverged) demonstrated by McKinney et al. (2017) with data simulations as well as empirical analyses, we fixed different cutoff value for each parameters diplayed (detailed of the cut-off values are reported in table S2). Black, red, blue and orange points represent singletons, duplicated, diverged and low confidence SNPs, respectively.

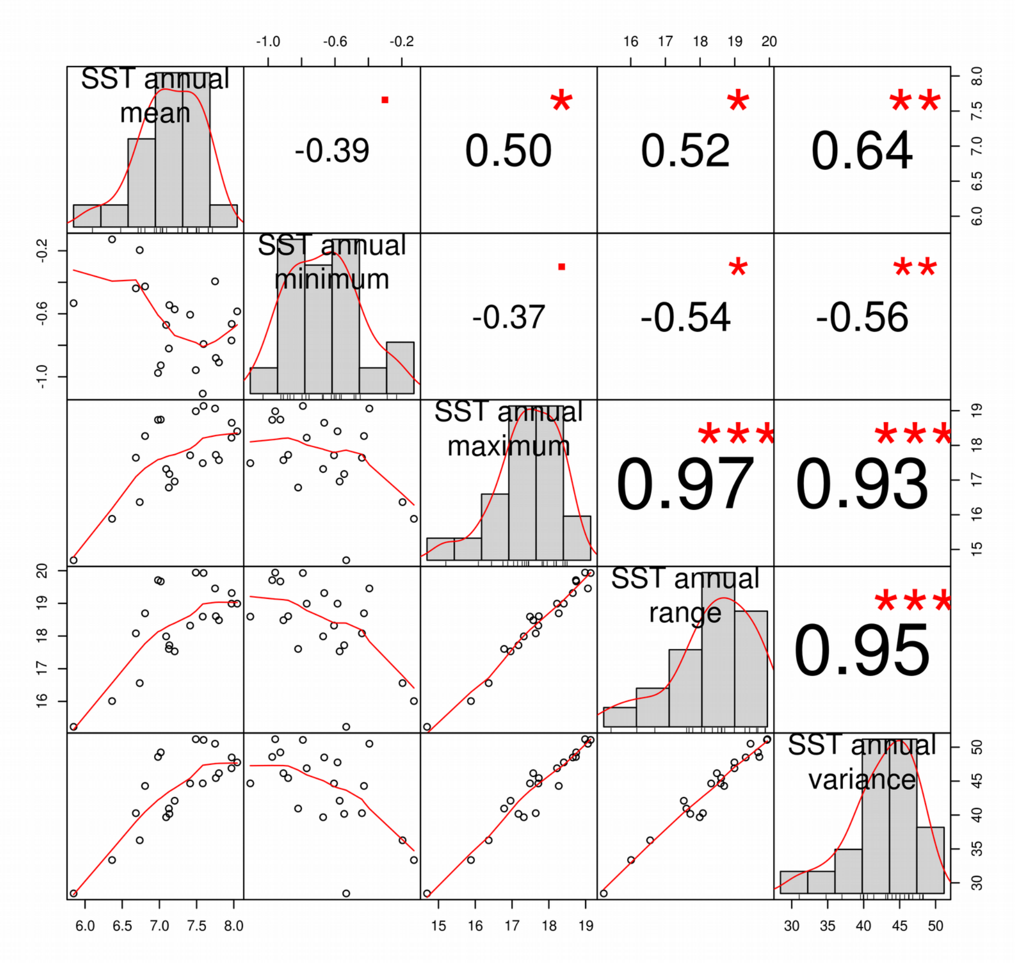


**Figure S2: Draftsman's plot with correlation among SST temperature parameters from the 21 sampling sites.** The distribution of each variable is shown on the diagonal. Below the main diagonal, the bivariate scatter plots with a fitted line are displayed. Above the main diagonal, the value correspond to Pearson's correlation coefficient plus the significance level as stars (i.e. p-values (0, 0.001, 0.01, 0.05, 0.1, 1) ; symbols("***", "**", "*"," "." ").

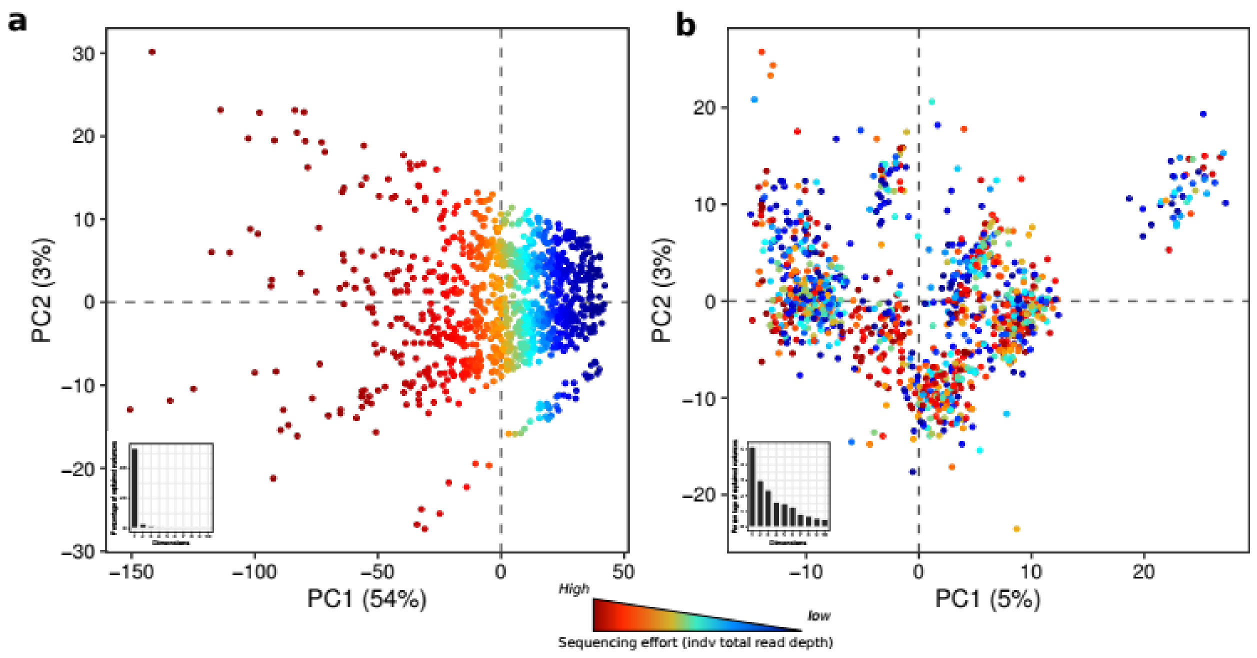


**Figure S3: Impact of read depth normalization for CNVs data**

**a.** PCA of individuals based on CNV loci read depth (not normalized); **b.** PCA of individuals based on CNV loci read depth normalized using the TMM method (Robinson & Oshlack, 2010). The color scale from blue to red represent the scale of sequencing effort (i.e. total number of reads sequenced for a given individual)

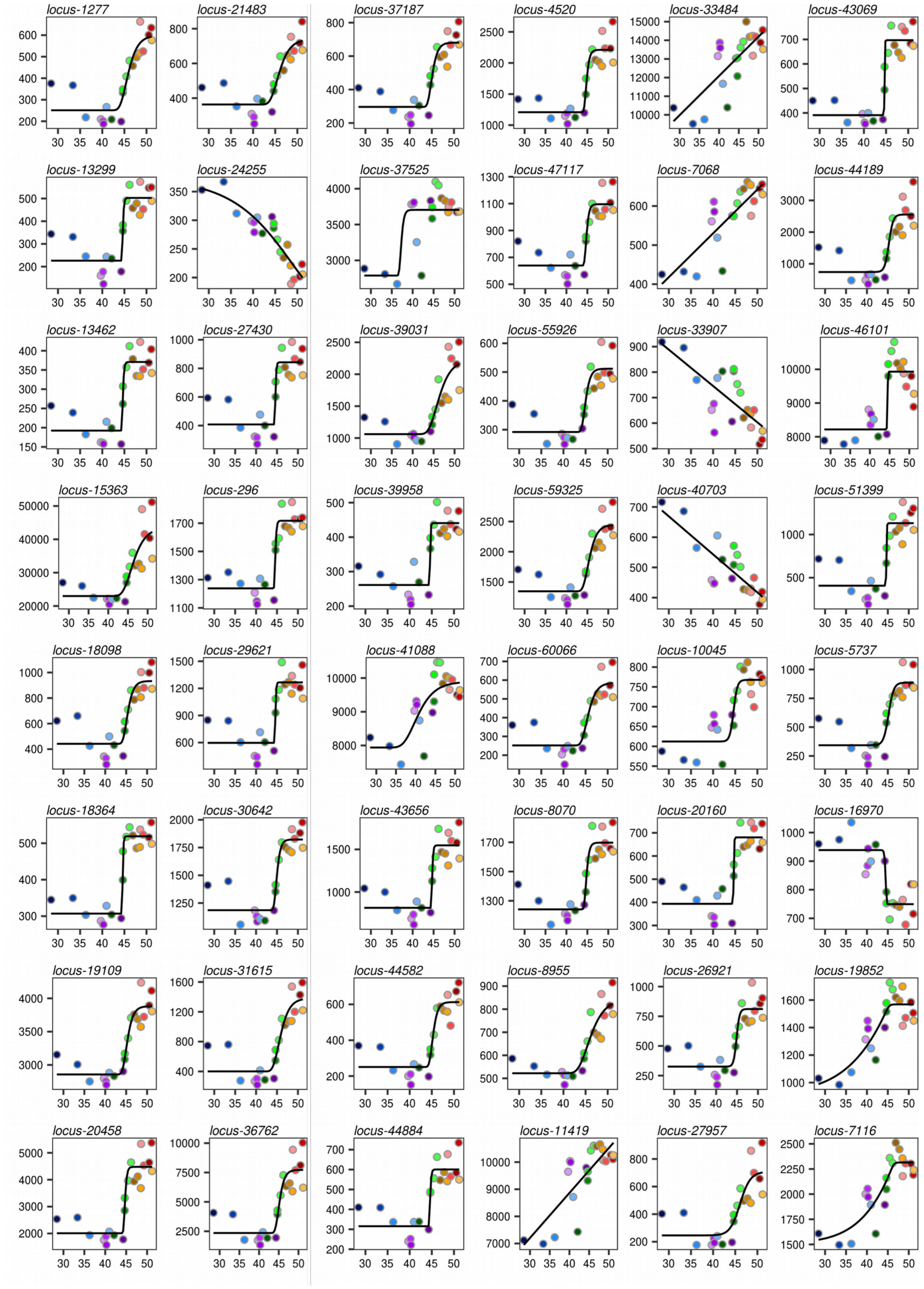


**Figure S4: Relationship between SST annual variance and CNVs read depth**

Scatter plot representing the relationship between SST annual variance (x-axis) and locus read depth (i.e. mean normalized read depth per sampling site; y-axis) for each of the 48 CNVs loci identified by both RDA and GLMM models. Each point corresponds to a sampling site colored according to the color code displayed in the figure 1a for each sampling site. The black line represents the best explicative model (see figure.S5 for details).

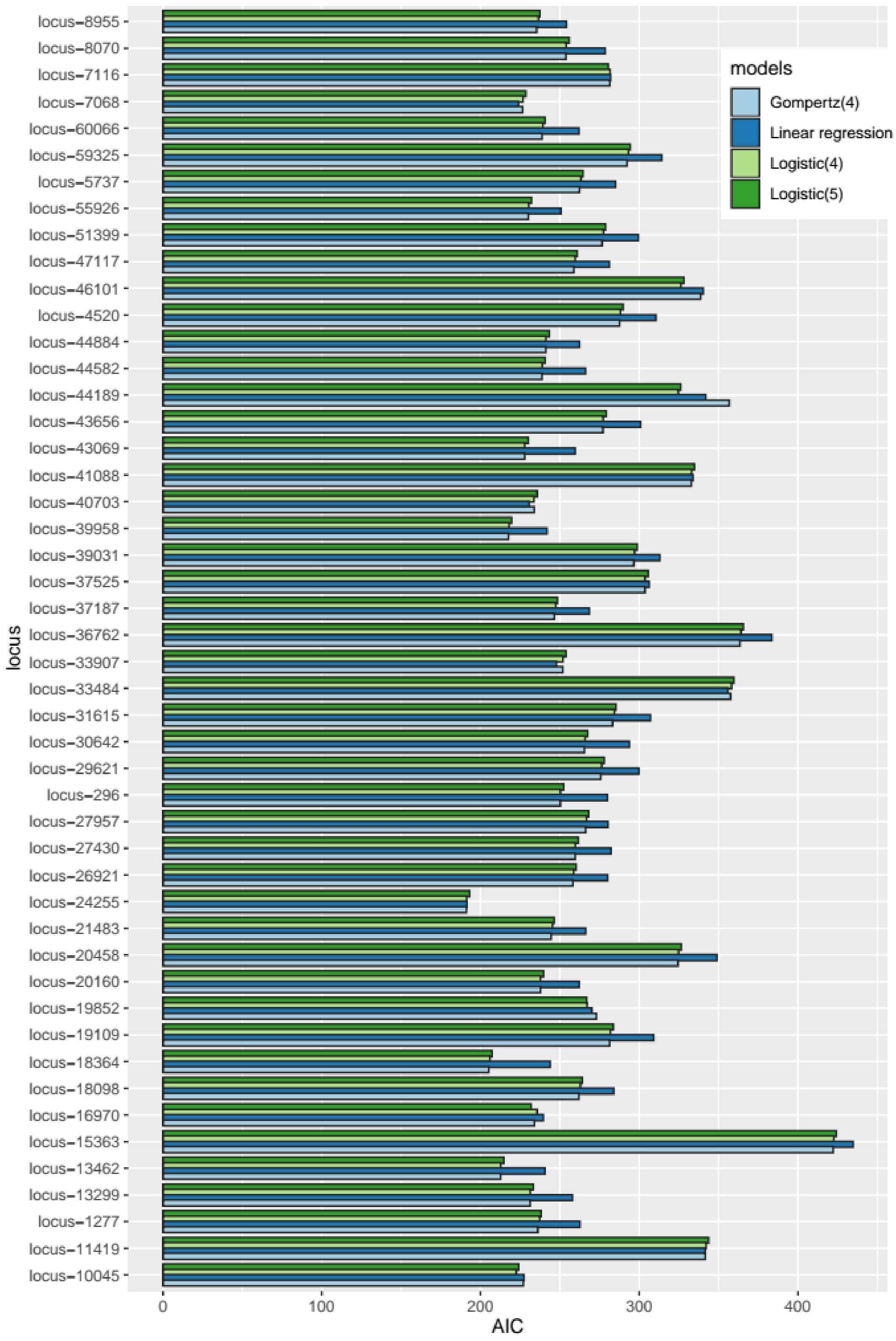


**Figure S5: Model comparisons**. Barplot showing the AIC coefficient of four different models (i.e. Gompertz with 4 parameters, logistic regression with 4 and 5 parameters and linear regression) assessing the relationship between read depth (i.e. mean normalized read depth per sampling site) and SST annual variance among 48 CNV loci associated with SST annual variance (i.e RDA and GLMM results).

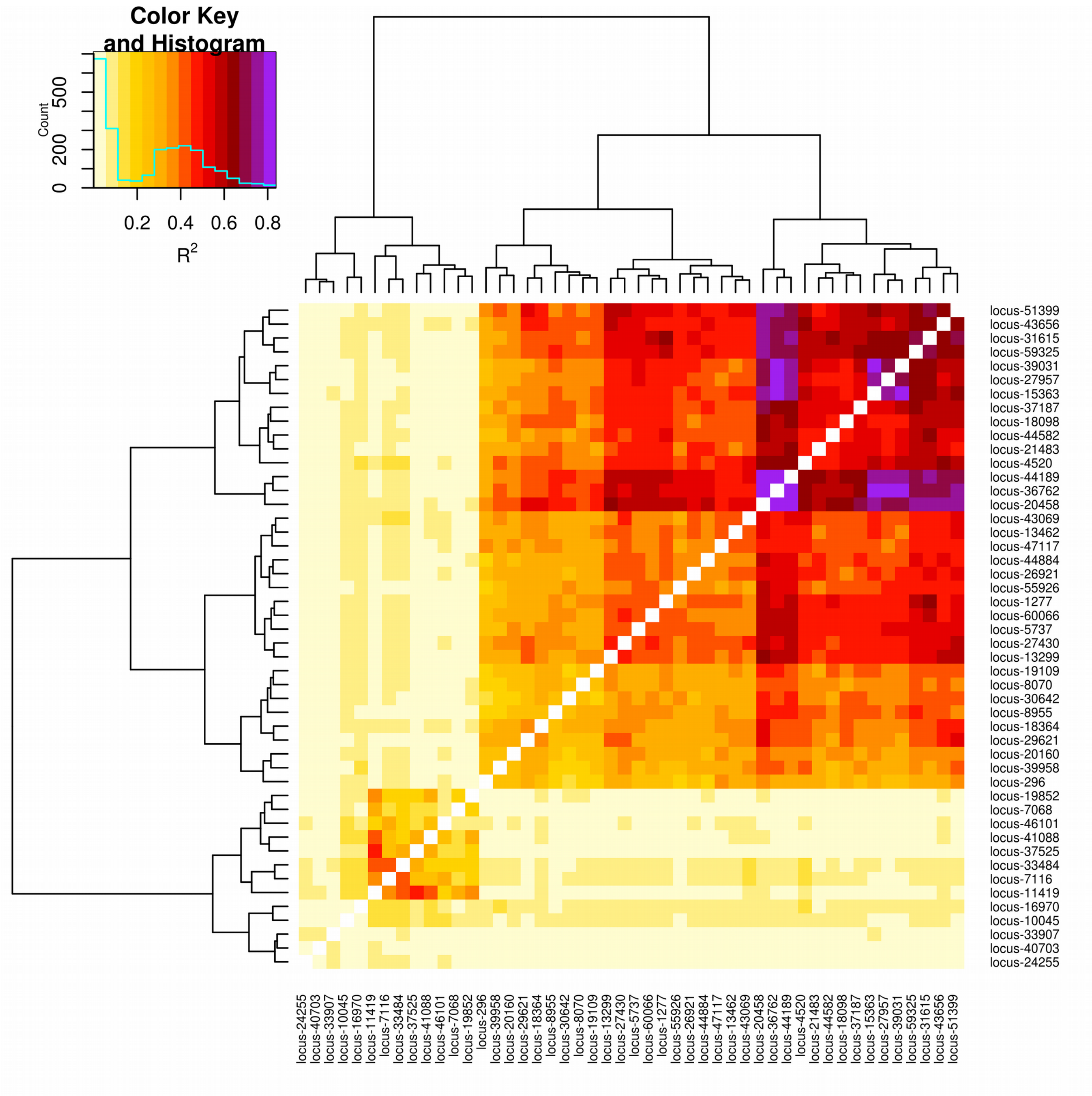


**Figure S6: Heatmap of statistical relationship among the 48 CNV loci associated with the SST annual variance.** Each cell represents the coefficient of determination (i.e. $R^2$) between each distribution of individuals' normalized read depth for a given pair of CNV loci. Loci are sorted based on an UPGMA clustering algorithm.

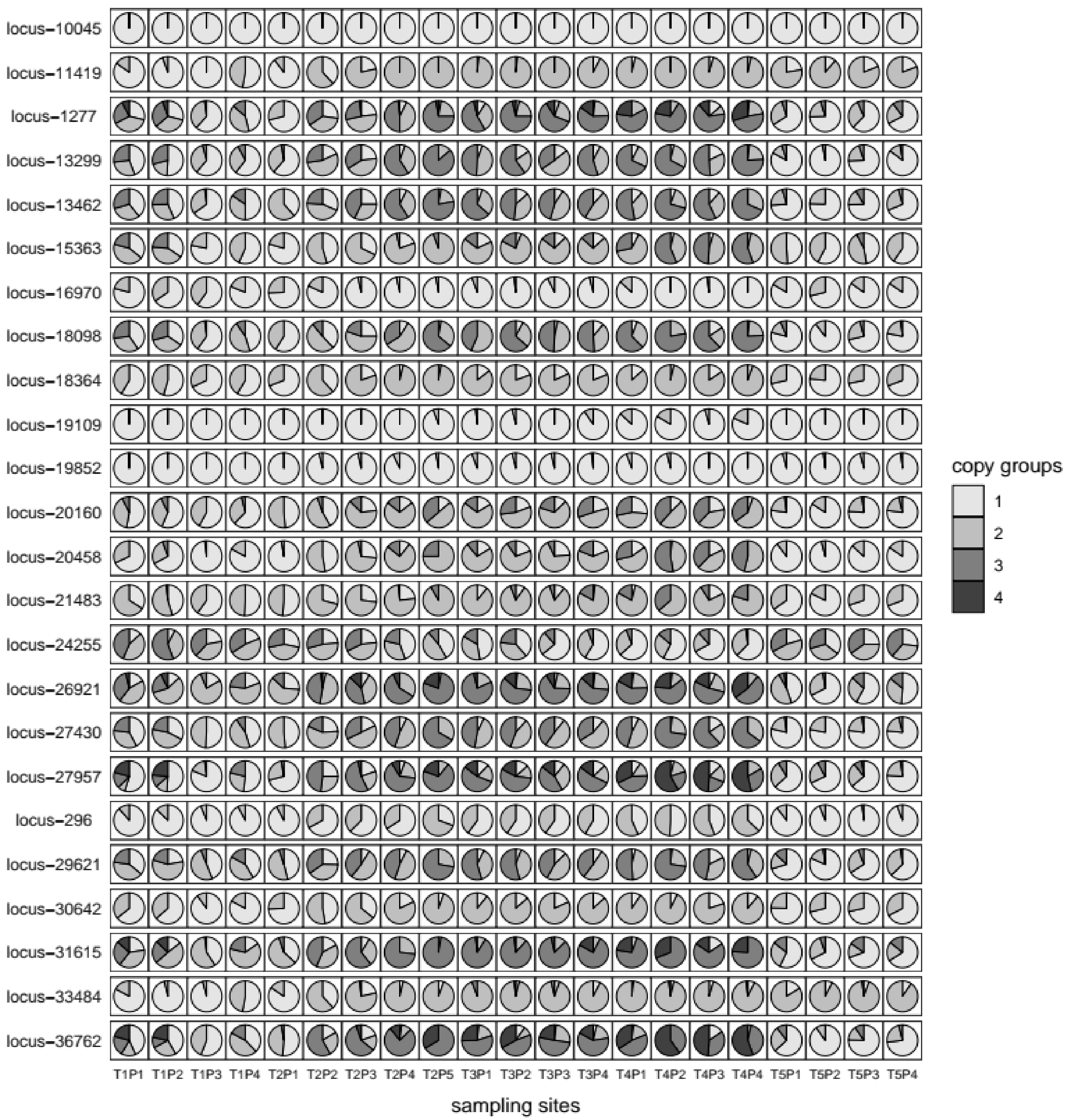

locus-10045
locus-11419
locus-1277
locus-13299
locus-13462
locus-15363
locus-16970
locus-18098
locus-18364
locus-19109
locus-19852
locus-20160
locus-20458
locus-21483
locus-24255
locus-26921
locus-27430
locus-27957
locus-296
locus-29621
locus-30642
locus-31615
locus-33484
locus-36762
T1P1 T1P2 T1P3 T1P4 T2P1 T2P2 T2P3 T2P4 T2P5 T3P1 T3P2 T3P3 T3P4 T4P1 T4P2 T4P3 T4P4 T5P1 T5P2 T5P3 T5P4
sampling sites
copy groups
1
2
3
4

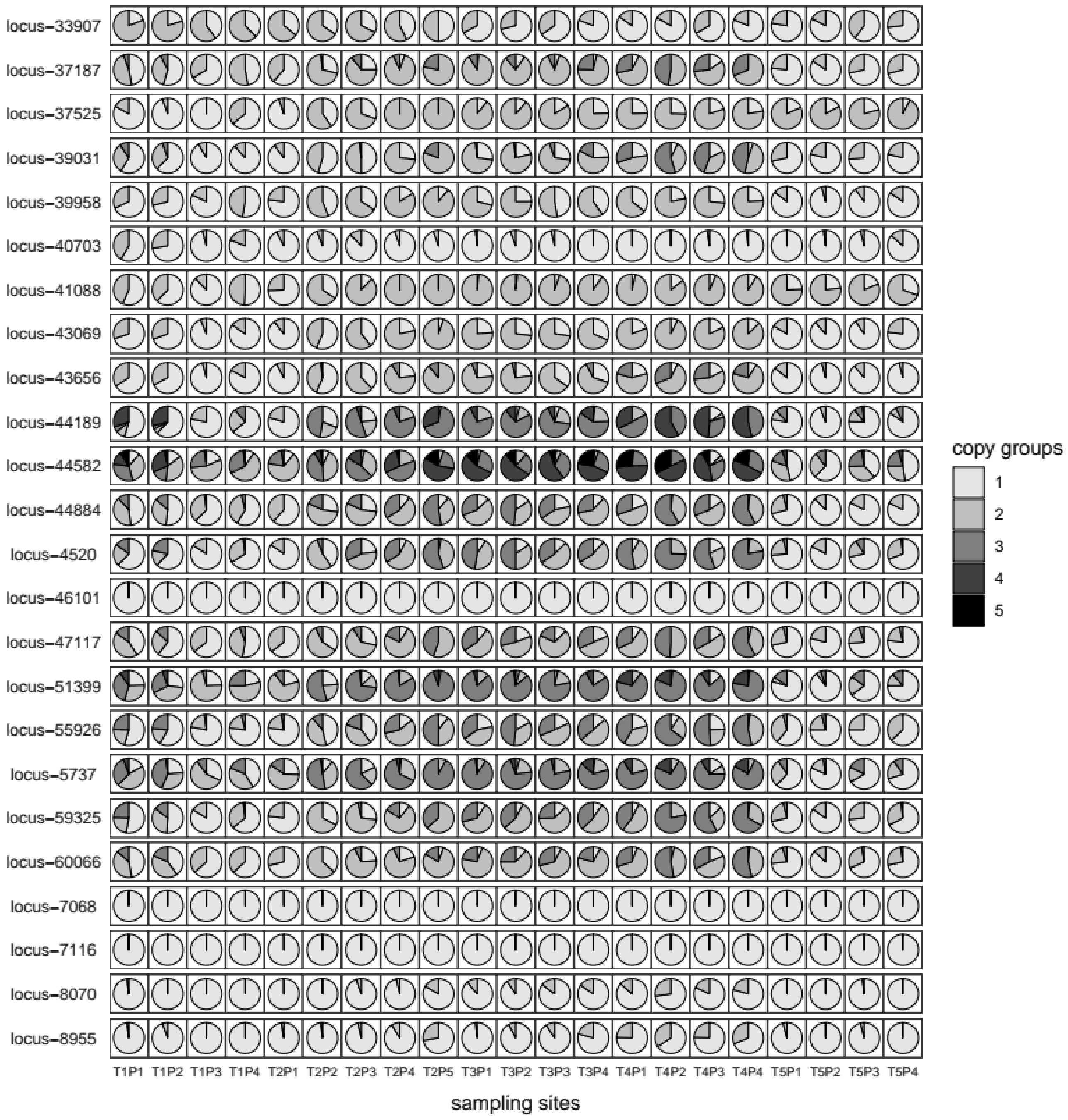


**Figure S7: Discretization of copy groups for 48 CNVs associated with SST annual variance.** Each pie chart represents the proportion of individuals classified within each 'copy group' among each sampling sites. Sampling sites are distributed along columns, CNV loci associated with annual variance of SST are distributed by rows (i.e. 48 loci that overlapped RDA and LME results). For each locus, the number of "copy groups" was estimated based on independent EM algorithm (see M&M for details).

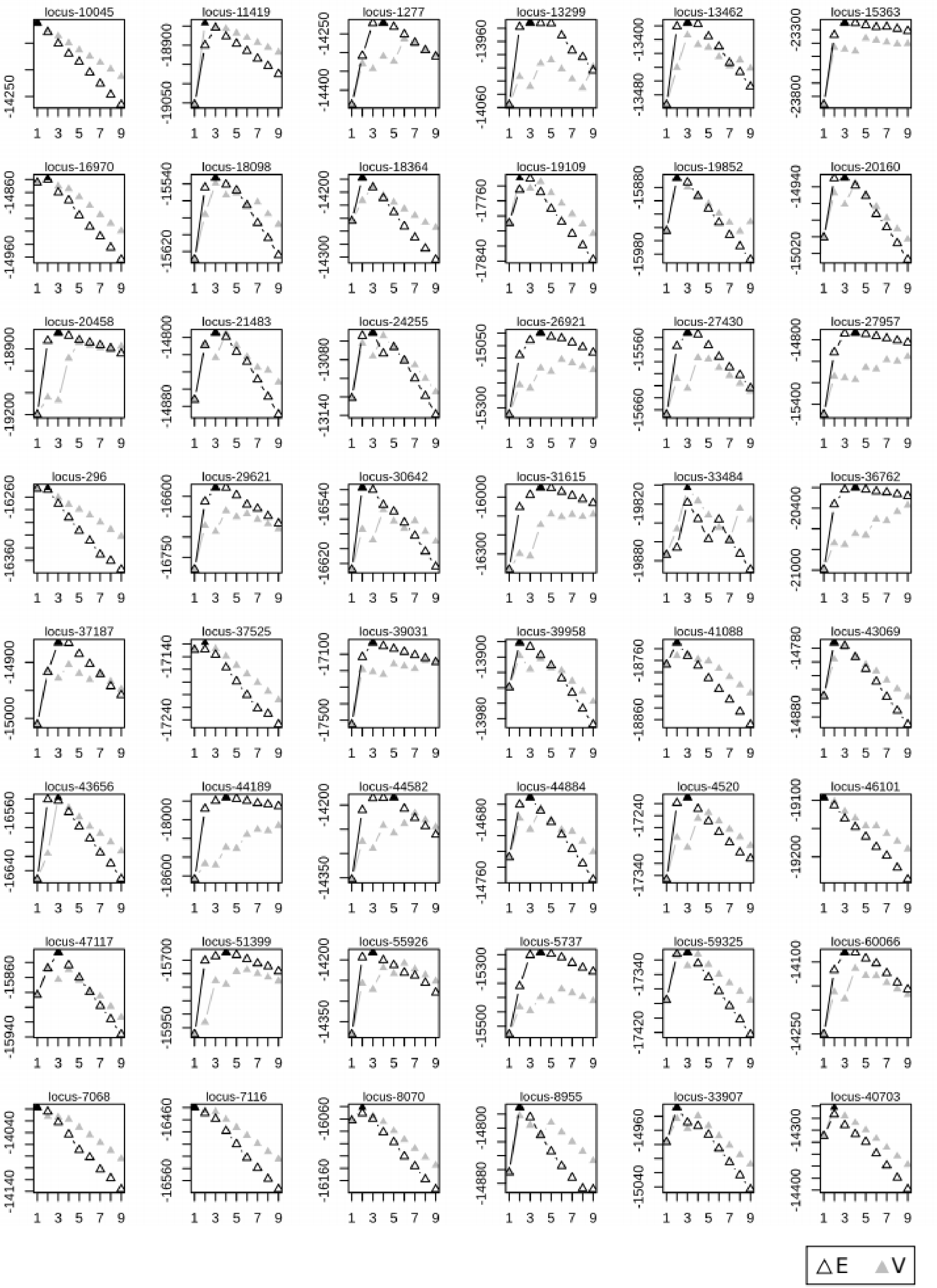


**Figure S8: Mclust model performance among the 48 CNV loci associated with SST annual variance.** (E) represents EM algorithm for equal variance (one-dim) and (V) represents EM algorithm for variable/unequal variance (one-dim). The black triangle in each figure illustrates the best model for sample clustering via the BIC (BIC = Bayesian Information Criterion) for up to 9 components.

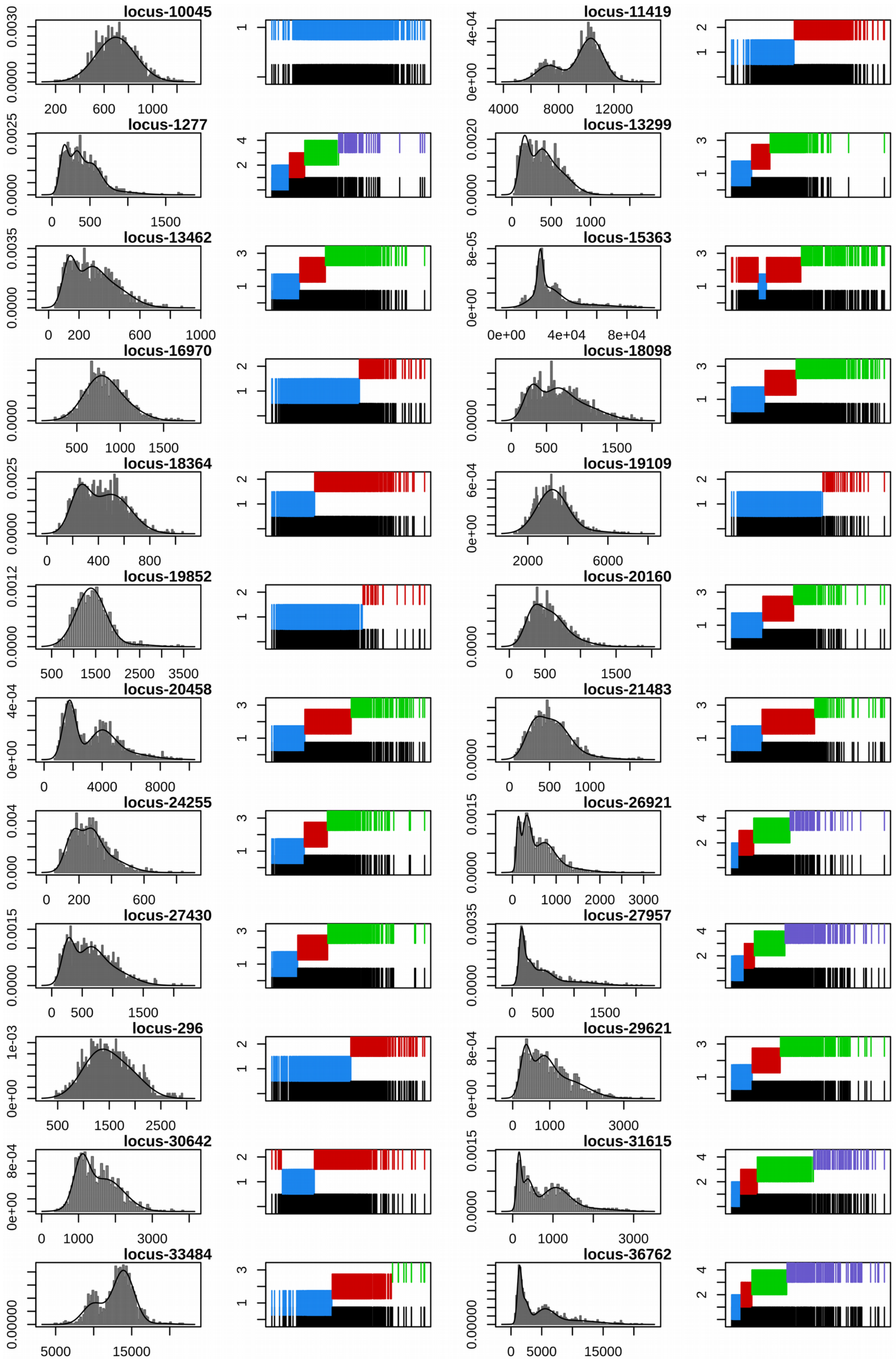

locus-10045
0.0000 0.0030
200 600 1000
locus-11419
0e+00 4e-04
4000 8000 12000
locus-1277
0.0000 0.0025
0 500 1500
locus-13299
0.0000 0.0020
0 500 1000
locus-13462
0.0000 0.0035
0 200 600 1000
locus-15363
0e+00 8e-05
0e+00 4e+04 8e+04
locus-16970
0.0000
500 1000 1500
locus-18098
0.0000
0 500 1500
locus-18364
0.0000 0.0025
0 400 800
locus-19109
0e+00 6e-04
2000 6000
locus-19852
0.0000 0.0012
500 1500 2500 3500
locus-20160
0.0000
0 500 1500
locus-20458
0e+00 4e-04
0 4000 8000
locus-21483
0.0000
0 500 1000
locus-24255
0.000 0.004
0 200 600
locus-26921
0.0000 0.0015
0 1000 2000 3000
locus-27430
0.0000 0.0015
0 500 1500
locus-27957
0.0000 0.0035
0 500 1500
locus-296
0e+00 1e-03
500 1500 2500
locus-29621
0e+00 8e-04
0 1000 3000
locus-30642
0e+00 8e-04
0 1000 3000
locus-31615
0.0000 0.0015
0 1000 3000
locus-33484
0.00000
5000 15000
locus-36762
0.00000
0 5000 15000

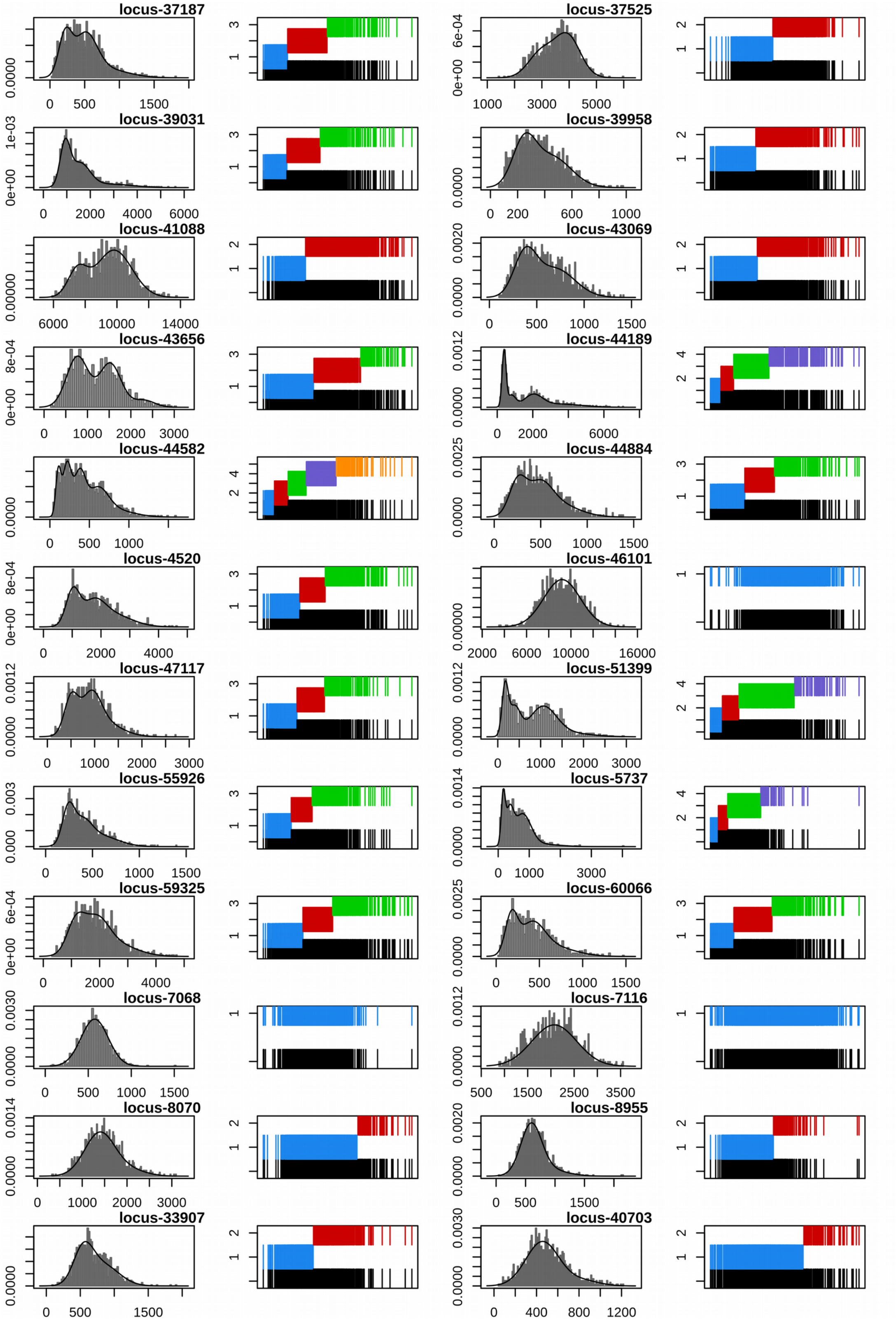


**Figure S9: Mclust model features for the 48 CNV loci associated with SST annual variance.** (left) Density of normalized read depth distribution (i.e. one-dimensional model); (right) classification plot from the best model (i.e. evaluated by BIC). In the classification plot, all individuals are displayed at the bottom (black bars), with the separated classes shown different levels above (colored bars).
